# Drosophila learn to prefer immobile spherical objects through repeated physical interaction

**DOI:** 10.1101/2025.06.09.658381

**Authors:** Kenichi Iwasaki, Sena Kawano, Ashnu Cassod, Charles Neuhauser, Aleksandr Rayshubskiy

## Abstract

Animals interact with unfamiliar objects to learn about their properties and guide future behavior, but the underlying neurobiological mechanism is not well understood. Here, we developed a behavioral paradigm in which freely walking *Drosophila melanogaster* are repeatedly guided to spherical objects using a visual cue. Flies exhibited diverse and structured object interaction motifs, including “ball pulling”, and “ball walking”, that evolved over time. Notably, flies developed a strong preference for immobile over mobile spherical objects, despite their near identical appearance, suggesting they learn about the object’s stability through physical interaction. This preference was impaired by silencing specific hΔ neurons in the fan-shaped body, previously implicated in spatial navigation but not known to contribute to object interactions. Our results show that hΔ neurons also modulate object interaction motifs and fidelity of following visual guidance cues, pointing to a role in balancing goal-directed and exploratory behaviors. These findings establish *Drosophila* as a model for investigating how internal representations and multimodal feedback contribute to adaptive object interaction.

## Introduction

Across the animal kingdom, organisms must contend with physical challenges unique to their local environments. A key challenge many species face is deciding which objects, among a multitude of animate and inanimate elements in their surroundings, are safe to approach and which should be avoided. The ability to physically engage with unfamiliar objects (hereafter “object interactions”) enables organisms to acquire information about an object’s properties and is widespread across taxa (Glickman and Sroges 1966; Lobo et al. 2014; Christensen et al. 2021; Galpayage Dona et al. 2022). Many species exhibit an intrinsic bias toward exploring novel objects, a behavioral trait thought to facilitate learning and environmental assessment (Berlyne 1950; Ibañez et al. 2023; Kidd and Hayden 2015; Ogasawara et al. 2022). Through such interactions, animals can gather critical data, such as whether an object poses a threat or offers a benefit, that cannot be inferred from vision or scent alone (Galpayage Dona et al. 2022; Lambert et al. 2017). Once familiar with an object, organisms can then choose among context-appropriate behavioral responses that enhance survival or reproductive success (Lambert et al. 2021; Liu et al. 2009; Schrauf, Huber, and Visalberghi 2008). Indeed, object interactions underpin a wide range of adaptive behaviors, including foraging, courtship display, and nest construction building (Bird and Emery 2009; Fayet, Hansen, and Biro 2020; Kelley and Endler 2012; Laumer et al. 2017; McCoy et al. 2019; Mizuuchi et al. 2018). Despite their broad ecological relevance, however, the neural mechanisms that regulate how animals engage with novel objects remain poorly understood.

The common fruit fly (*Drosophila melanogaster*) is a leading model for investigating the biological basis of complex behaviors, owing to its extensive genetic toolkit and accessibility. Much of what we know about *Drosophila* complex behavior comes from studies of social interactions such as courtship, aggression, and group dynamics (Alekseyenko et al. 2014; Billeter et al. 2009; Coen et al. 2014; Deutsch et al. 2019; Hoopfer et al. 2015; Jiang et al. 2020; Schretter et al. 2020; Hindmarsh Sten et al. 2025; Sun et al. 2020; Wang et al. 2011; Dukas 2020). In contrast, far less is understood about how flies interact with the physical environment beyond the social context. Particularly, how they physically engage with objects, whether they can learn about physical properties of objects, and how such learning may influence future decisions.

Flies are known to form associations between spatial locations and sensory cues, including visual (Ofstad, Zuker, and Reiser 2011) and gustatory stimuli (Corfas, Sharma, and Dickinson 2019; Kim and Dickinson 2017; Behbahani et al. 2021). However, it remains unclear whether similar associations extend to the physical attributes of objects themselves. During flight, flies tend to avoid small objects, presumably to minimize risk of collision or predation (Cheng, Colbath, and Frye 2019; Maimon, Straw, and Dickinson 2008; Mongeau et al. 2019). Interestingly, this aversion can be reversed when small objects are paired with attractive sensory cues (Cheng, Colbath, and Frye 2019). Flies also show a preference for perching on taller structures, possibly to gain visual vantage points for environmental assessment, mirroring behaviors seen in naturalistic settings (Robie, Straw, and Dickinson 2010). Together, these findings suggest that flies can flexibly evaluate and respond to object features in a context-dependent manner. However, whether and how such object-related decision-making occurs during terrestrial exploration remains largely unexplored. This gap is due, in part, to the difficulty of eliciting repeated, sustained interactions between freely walking flies and inanimate objects.

We previously demonstrated that the movement of freely walking flies can be precisely guided toward a spatial target using a dynamic visual cue (Iwasaki et al. 2025). In the present study, we extend this approach to direct flies toward discrete objects, enabling repeated and sustained interactions suitable for detailed behavioral analysis. Using this visual guidance paradigm, we found that male flies are capable of complex and structured object interactions with a small spherical target (ball). These interactions consisted of multiple distinct motor motifs, some involving coordinated, multi-step behaviors not previously described in *Drosophila*. Flies flexibly modulated their interactions based on object properties: they consistently preferred immobile over mobile balls and were able to discriminate between the two even when interacting with both repeatedly in the same session. To investigate whether circuits implicated in spatial navigation contribute to object preference, we examined the role of hΔ neurons in the central complex. While only the hΔB subtype has been directly linked to vector-based navigation (Hulse et al. 2021; Lyu, Abbott, and Maimon 2022), the functions of most hΔ subtypes, of which there are over a dozen, remain poorly understood, particularly in the context of object-directed behavior. Silencing specific hΔ subtypes disrupted the development of object preference and altered the expression of distinct motor motifs. These findings point to a previously unrecognized role for internal spatial computations, possibly involving vector arithmetic, in guiding object-directed interactions. Together, our results uncover new layers of physical and cognitive sophistication in *Drosophila* behavior and provide a framework for mechanistic studies of adaptive object engagement.

## Results

### Visually guided flies engage in persistent interactions with a spherical object

Previous work has shown that freely walking flies can be visually guided to specific spatial destinations using a rotating “pinwheel” cue, which dynamically adjusts its rotation based on the fly’s position and orientation (Iwasaki et al. 2025). To test whether this visual navigation system could promote interactions with discrete objects, we used it to repeatedly direct flies toward a small, smooth, plastic spherical object (2.38 mm in diameter, 10 mg mass; Figure 1A), which is similar in size but approximately 10 times heavier than the fly itself (body length ∼2.5 mm; mass ∼1 mg). The guidance cue was deactivated once the fly approached within a 5-mm radius of the object, allowing the fly to freely decide whether to interact (Figure 1B). The guidance cue was reactivated once the fly moved beyond a 5-mm radius from the object. Flies were not required to engage with the object and could leave without penalty.

**Figure 1.**
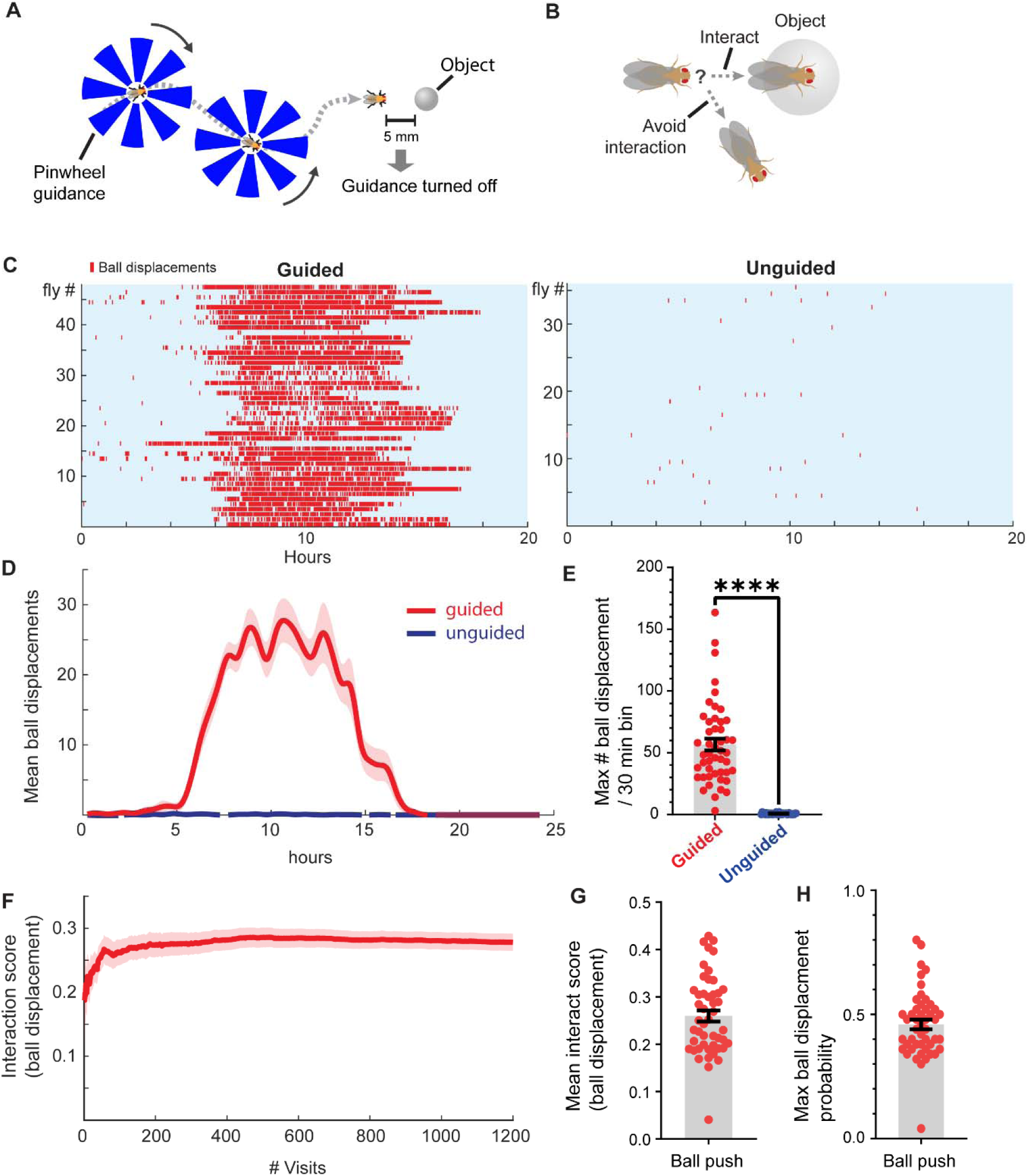
Visual guidance enables repeated fly-object interactions with novel spherical targets. **(A)** In the “pinwheel-guided” assay, flies are directed toward a spherical object using a visual stimulus (the “pinwheel”). Guidance is turned off when the fly reaches within a 5-mm radius of the object and reactivated once the fly moves beyond the 5-mm radius. **(B)** The decision to engage with the object is made autonomously by the fly. **(C)** Guided flies frequently engage with and move the balls, while unguided flies rarely do so (*n* = 48 guided flies, 36 unguided flies). **(D)** Mean frequency of ball displacement events per 30-minute bins over time for guided and unguided flies (n = 48 guided, 36 unguided). The delayed onset of interactions (beginning approximately hour 5) parallels the rise in general locomotor activity observed in Figure S1E. **(E)** Maximum ball displacement rate (displacement events per 30-minute bin) for each group. *Mann–Whitney test*, two-tailed (*n* = 48 guided flies, 36 unguided flies). **(F)** Cumulative interaction scores (ball displacements) over number of visits. Only flies with >1,200 visits included (*n* = 32 flies). The >1200 push threshold was chosen to highlight a subset of flies that engaged in a particularly high number of ball interactions, thereby demonstrating the sustained and repeated nature of the behavior. **(G)** Mean interaction score (ball displacement events per visit) across flies (*n* = 48 flies). **(H)** Maximum ball displacement probability achieved per fly. Probability was calculated over a sliding window of 50 visits, and the highest value recorded for each fly was used (*n* = 48 flies).

We found that most guided flies initiated active, repeated interactions with the ball, often displacing it across the arena over extended periods (Figures 1C–1D). By contrast, flies that were not visually guided interacted with the ball only rarely. Guided flies moved the ball on approximately 26% of their visits, with some reaching a maximum ball-displacement probability of approximately 50% (Figures 1F–1H).

To test whether the increase in object interaction could be explained by passive exposure to the visual cue or simple proximity to the ball, we conducted two control experiments. One possibility was that the presence of the visual stimulus itself might enhance the salience of the nearby ball, thereby increasing the likelihood of interaction. First, presenting stationary (non-rotating) pinwheels (centered either on the fly or on the ball) did not increase interaction frequency, although it slightly reduced average fly-to-ball distance (Figures S1A, S1B). Second, reducing the arena size to force proximity between flies and balls failed to produce comparable levels of interaction (Figures S1C, S1D). These results indicate that dynamic, trajectory-coupled visual guidance, not mere pinwheel presence or spatial closeness, is critical for promoting sustained fly-ball interactions.

### Guided flies exhibit a repertoire of distinct interaction motifs with a spherical object

During initial experiments, we observed that guided flies interacted with the ball using a range of distinct strategies (Videos 1–9). These included walking across the ball without displacing it, displacing the ball while jumping off it, and pulling or displacing the ball from the side while walking on the ball. These observations suggested that flies engage with the object through a variety of structured behaviors. To quantify this diversity, we measured three kinematic features for each interaction: ball displacement during interaction (before the fly exited the ball), ball displacement after interaction (after exit), and fly movement after interaction. Fly movement after interaction was analyzed over a 200-millisecond window, based on the finding that most jump events occurred within this period (Figures S2A, S2B).

We applied k-means clustering to 29,982 interaction events collected from 48 male flies, identifying five distinct behavioral motifs (Figure 2A). These included “walk-off” events, in which flies crossed over the ball without substantial movement of either the fly or the ball; “ball pull” events, where the fly displaced the ball while dismounting with minimal post-interaction fly movement; and “ball walk” events, characterized by the fly walking along the side of the ball while rolling it. Two additional clusters captured jump-like behaviors, defined by rapid post-interaction fly movement with varying degrees of ball displacement (Figure 2B). Importantly, these interaction motifs were not restricted to particular individuals: most flies contributed to all five clusters, with no single fly accounting for more than 12% of the events in any one group (Figure S2C).

**Figure 2.**
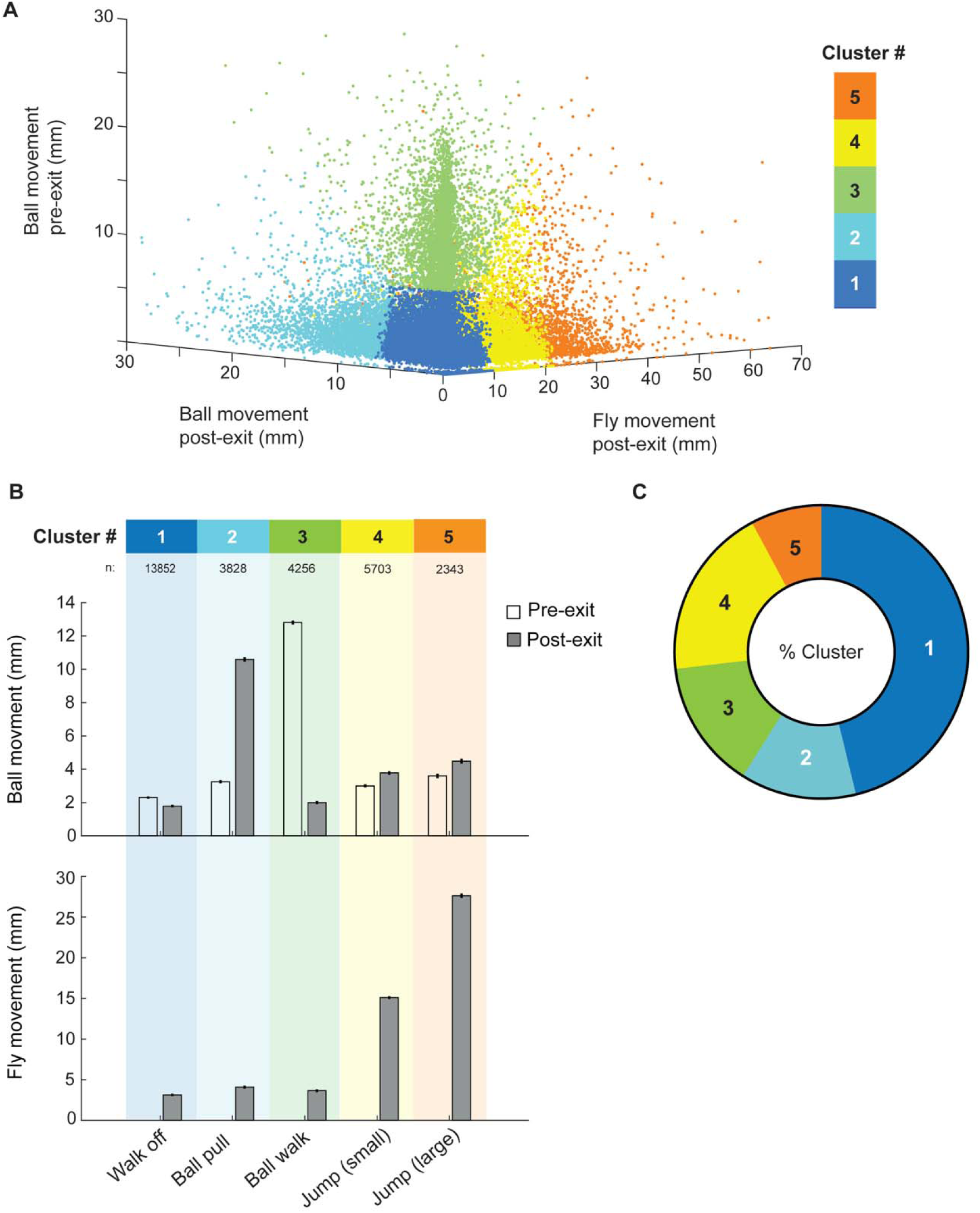
Unsupervised clustering reveals distinct categories of fly-ball interaction motifs. **(A)** K-means clustering was applied to 29,982 interactive events from 48 male flies, using features related to ball displacement (before fly exit (pre-exit) and after fly exit (post-exit)) and fly movement following interaction termination (post-exit fly displacement). A five-cluster solution was chosen to isolate “walk-off” events – characterized by minimal ball and fly movement – from other motif types while minimizing redundancy. **(B)** The clustering analysis revealed several distinct behavioral motifs, including “walk-off,” “ball pull,” “ball walk,” and “jump”-like events. **(C)** Among all classified events, “walk-off” interactions were the most frequent (46%), followed by small jumps (19%), ball walks (14%), ball pulls (13%), and large jumps (8%).

### Interactive motifs follow distinct patterns of temporal development

Having identified multiple fly-ball interaction motifs through clustering (Figure 2), we next asked whether these motifs were used consistently throughout the experiment or evolved differently over time. Divergence in their temporal trajectories would suggest that different interaction strategies may be governed by distinct regulatory processes in the nervous system.

To address this, we focused on two distinct behavioral motifs: “ball pull” and “ball walk.” Ball-pull events were defined by displacement of the ball after the fly departed, whereas ball-walk events involved the fly actively moving the ball while walking along its side (Figure 3Ai–iv). Full definitions and classification criteria for these motifs are detailed in the Methods. These two behaviors exhibited markedly different temporal profiles. Ball-walk frequency gradually increased over the course of the experiment, peaking in the later hours, whereas ball-pull frequency rose initially but plateaued early (Figure 3B). This divergence was confirmed both in raw event counts and in normalized comparisons of early and late time bins: flies performed significantly more ball walks in the second half of the session, while the frequency of ball pulls remained unchanged (Figures 3C, 3D; see also Figures S2D, S2E).

**Figure 3:**
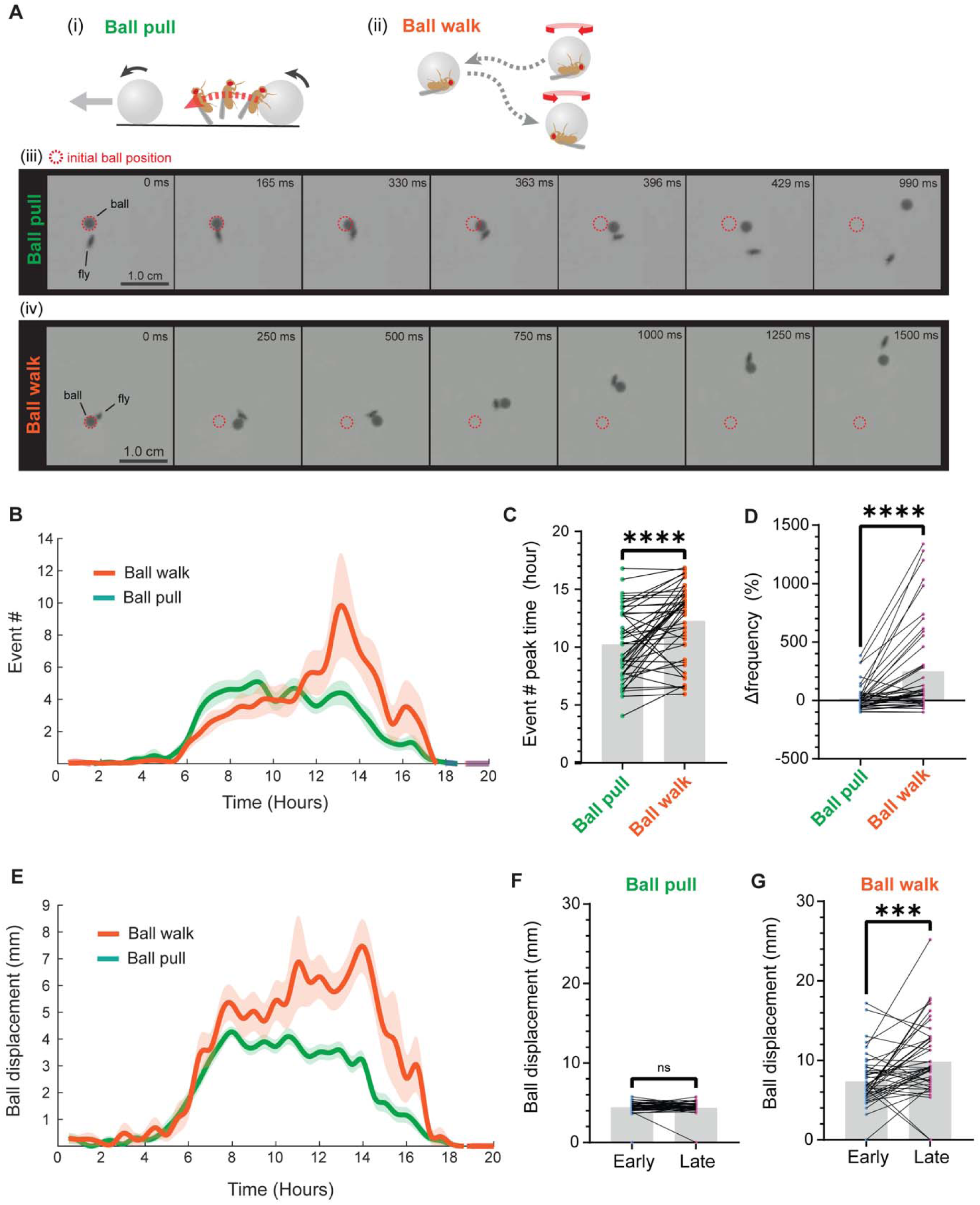
Ball-walk interactions show distinct temporal development from ball-pull events. **(A)** Schematic examples of (i, iii) *ball pull* and (ii, iv) *ball walk* events. In ball pull, a fly mounts the ball and pulls it upon dismounting, resulting in a displacement (≥3 mm) without walking on the ball. In ball walk, a fly remains on the side of the ball and displaces it by walking along its surface; these events are defined by coordinated fly-ball movement (>2.5 mm). Definitions are based on movement thresholds exceeding noise levels (<2.5 mm). The red dotted circles indicate the ball’s original position. In ball walk events, significant displacement occurs while the fly remains on the ball, suggesting active control during locomotion. In contrast, ball pull events are defined by displacement that occurs after the fly dismounts, as the fly appears to “throw” the ball upon exit. **(B)** Time-course of motif frequencies across the experimental period (30-minute bins). Ball-walk frequency increases gradually and peaks later, whereas ball-pull frequency plateaus earlier. *Y*-axis: mean number of events; *X*-axis: experimental time in hours. *n* = 48. **(C)** Ball pull frequency peaks significantly earlier than ball walk frequency (*Wilcoxon matched-pairs signed-rank test, two-tailed*, *n* = 47 flies; one fly excluded due to no relevant events). **_(D)_** Relative frequency change over time. Ball-walk frequency shows a significantly greater increase from early to late periods than ball-pull frequency. Δ frequency (%) = [(# events _late period_ - # events _early period_)/# events _early period_] x 100. Periods were divided in half per fly. *Wilcoxon matched-pairs signed-rank test, two-tailed*, *n* = 48 flies. **(E)** Average ball displacement distances for ball-pull and ball-walk events. *n* = 48. **(F,G)** Ball displacement size comparisons (*Wilcoxon matched-pairs signed-rank test, two-tailed*) across early vs. late active periods for (F) ball pull (*n* = 48) and (G) ball walk (*n* = 48).

A cluster-based analysis confirmed the same pattern, independently supporting the conclusion that these motifs follow distinct temporal trajectories (Figures S2H, S2I). Notably, the distance the ball was displaced during ball walks increased significantly over time (Figure 3E,G), indicating that flies engaged in longer ball walks as the experiment progressed. In contrast, the magnitude of displacement during ball pulls remained stable (Figure 3F). These findings suggest that the increase in ball-walk frequency and displacement is not merely due to increased activity.

We also observed that the frequency of ball walks peaked as the frequency of jumps declined (Figures S2F, S2G), suggesting a possible shift in behavioral state. Together, these results indicate that interactive motifs are not only behaviorally distinct but also follow different patterns of temporal development, consistent with the idea that they may be under separate regulatory control.

### Flies develop preference for immobile spherical objects

In earlier experiments, the plastic balls used were fully mobile and could be readily displaced by most test flies. This raised a key question: how does object mobility influence fly behavior? Does the ability of the ball to move enhance interaction by providing sensorimotor feedback, or could it instead discourage engagement by making the object unpredictable or harder to evaluate? To address this, we directly tested how object mobility affects fly-ball interactions by comparing fly behavior toward two visually identical balls – one free to move and the other fixed in place using a small amount of glue. To control for potential olfactory differences arising from the glue, we blocked olfactory input by covering both antennae with UV glue (Figure 4A).

**Figure 4.**
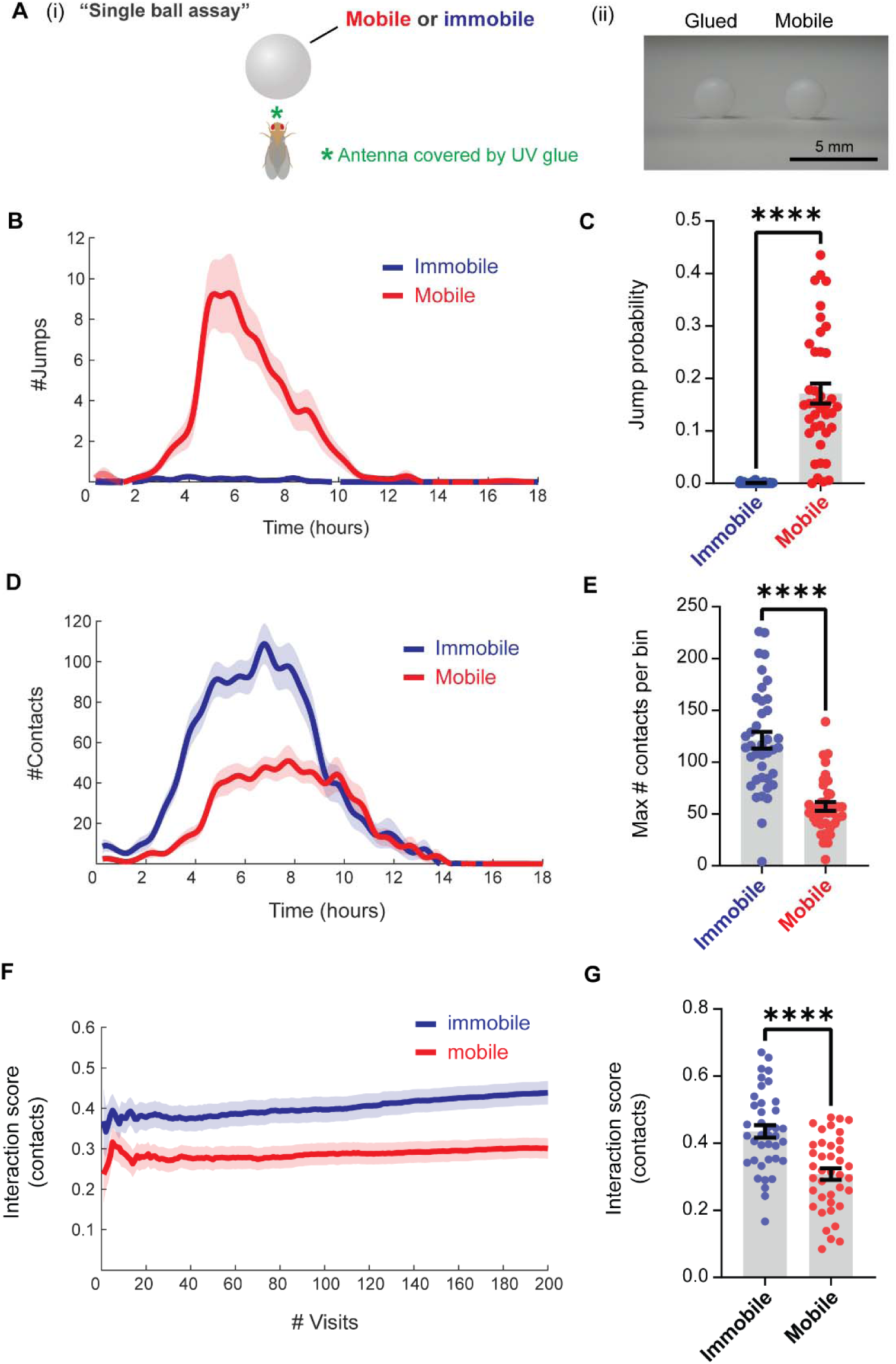
Flies preferentially interact with immobilized balls. To examine how ball mobility affects interaction behavior, we compared responses to mobile versus immobilized balls in a single-ball assay using flies with UV-glued antennae to block olfactory sensation. **(A)** (i) Schematic of the single-ball assay. Flies were guided toward either a mobile or glued (immobile) ball. (ii) A photograph of the glued (left) and mobile (right) balls, which are visually indistinguishable. **(B)** Flies interacting with mobile balls exhibited a high number of jump events (>5 mm displacement), whereas such jumps were rare when flies interacted with glued balls (*n* = 39 flies per group; 30-minute bins). The y-axis indicates the number of events per 30-min time bin. **(C)** The probability of jumping off a ball was significantly higher when the ball was mobile, indicating that mobility influences behavioral outcomes upon dismount. *Mann-Whitney Test*, two-tailed (*n* = 39 flies per group). **(D)** Contact frequency over time for mobile versus immobile balls (*n* = 39 flies per group). The y-axis indicates the number of events per 30-min time bin. **(E)** The maximum number of contacts per time bin was significantly higher for immobilized balls (*Mann–Whitney test*, two-tailed; *n* = 39 flies per group). **(F)** Interaction scores (contact) across the first 200 visits (only flies with ≥200 visits included; *n* = 38 flies per group). **(G)** Global interaction scores (total # contacts/ total # visits) were significantly higher for immobile balls. That is, flies were more likely to contact an immobilized ball than a mobile one. Final interaction scores across the entire experiment were used for statistical comparison to avoid bias from arbitrary visit windows (*unpaired t-test*, two-tailed; *n* = 39 flies per group).

Flies responded differently to the two ball types. Interactions with the mobile ball were marked by frequent jumping, whereas flies rarely jumped from the immobile ball (Figure 4B,C). Contact events, where the fly touched the ball, were significantly more common for the immobile ball (Figure 4D,E). Notably, the decision to contact the ball was made autonomously by the fly and likely reflects a combination of visual and mechanosensory evaluation during close-range inspection, shaped by prior experience.

Flies displayed a strong and persistent preference for the immobile ball, as reflected in their interaction scores across visits (Figure 4F,G). This preference was also observed in flies with an intact olfactory system (Figure S3) and remained significant when directly compared across conditions (Figure S4E). Interestingly, flies with functional olfaction exhibited even higher contact interaction scores (Figure S4F), suggesting that antennal olfactory or mechanosensory cues, or a combination of both, may enhance fly–object interactions.

Finally, flies spent significantly more time not moving (halting) beside immobile balls (Figures S4A-S4D), providing further evidence that they are capable of distinguishing physical object properties through direct interaction.

### Flies’ preference for immobile spherical objects persists even when they interact with two objects in the same session

The experiments described above involved a single object per experiment. To examine whether flies could discriminate between objects presented in parallel, we designed a two-object assay in which one ball was immobile and the other mobile. Flies were guided alternately between the two balls using our visual pinwheel cue, with each target switched only after the fly had completed an interaction with the current ball (Figure 5A). To eliminate the influence of olfactory cues on preference, we UV-glued the antennae in all experiments.

**Figure 5.**
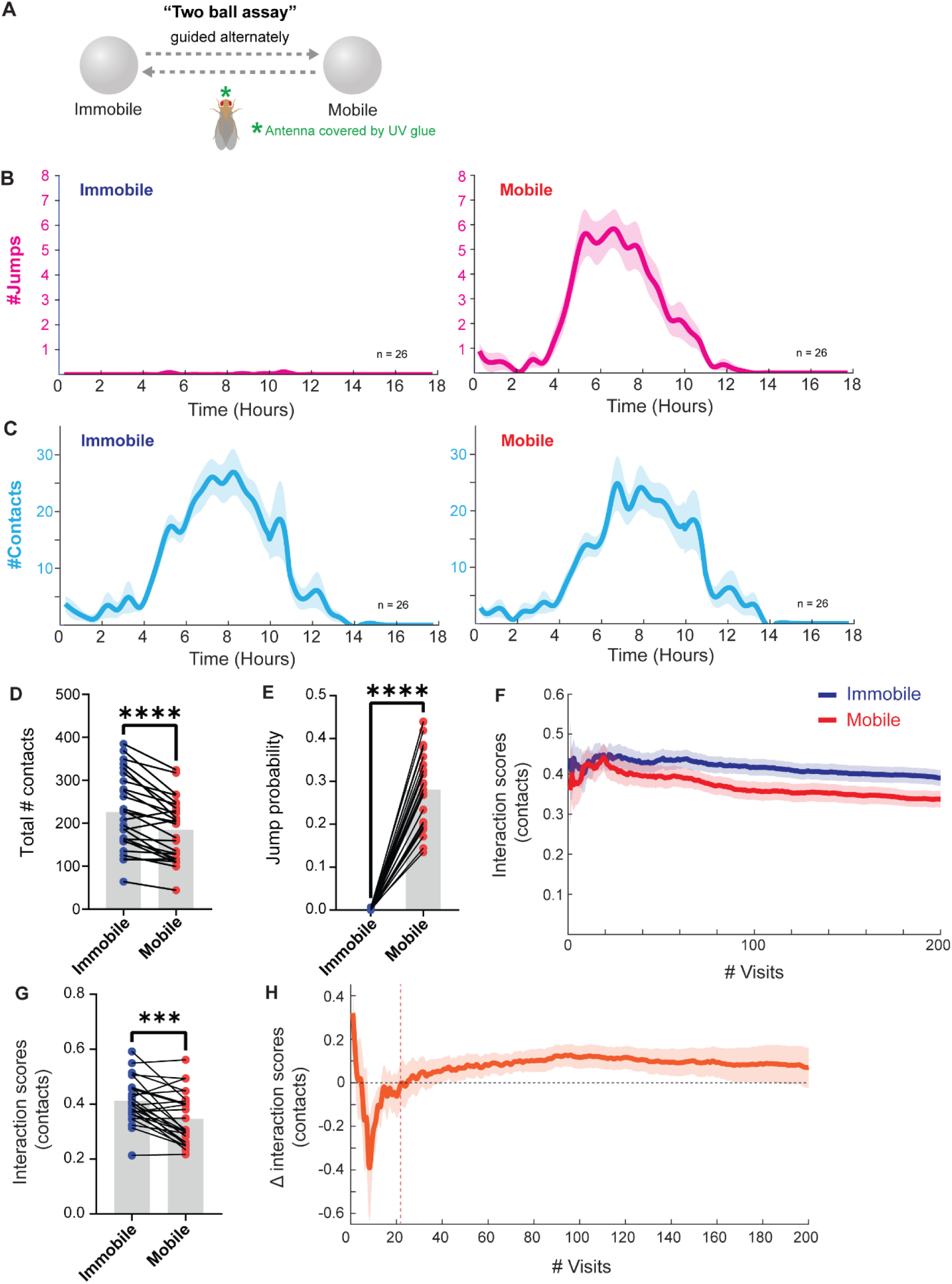
Flies develop a preference for immobilized balls when alternately interacting with mobile and immobile objects. To test whether flies distinguish between objects based on mobility alone, we designed a two-ball assay in which one ball was mobile and the other was immobile. Flies were guided to alternate between the two. Antennae were UV-glued to block olfactory input. **(A)** Schematic of the two-ball assay: flies alternately approached a glued (immobile) and a free (mobile) ball within the same experimental session. **(B)** Time course of jumping events for immobile (left) and mobile (right) balls (*n* = 26 flies per condition; 30-minute bins). The y-axis indicates the number of events per 30-min time bin. **(C)** Time course of contact events for immobile (left) and mobile (right) balls (*n* = 26 flies per condition; 30-minute bins). All jump events are a subset of contact events, each jump is preceded by physical contact with the ball. **(D)** Total number of contacts was significantly higher for immobile balls (*paired t-test*, two-tailed; *n* = 26 flies). **(E)** The probability of jumping after ball interaction was significantly higher for mobile balls (*Wilcoxon matched-pairs signed-rank test*, two-tailed; *n* = 26 flies). **(F)** Interaction scores (contacts) for mobile and immobile balls during the 1^st^ 200 visits (*n* = 25 flies). **(G)** Interaction scores (contacts) were significantly higher for immobilized balls (*paired t-test*, two-tailed; *n* = 26 flies). **(H)** Difference in interaction (contact) scores (Δ interaction score = score _immobile_ – score _mobile_) plotted as a function of visit number. A stable preference for immobile balls emerged after approximately 20 visits on average, following an initial dip (*n* = 25 flies).

As in the single-ball experiments (Figure 4), flies showed a clear behavioral distinction between the two object types. They frequently jumped off mobile balls but rarely from immobile ones (Figures 5B, 5E). Ball contact events occurred more often with immobile balls (Figures 5C, 5D), and this difference was consistent across experiments, yielding a persistent preference for immobile objects (Figures 5F, 5G). This preference stabilized after approximately 40 visits (Figure 5H), suggesting that repeated exposure supports the formation of a stable preference. Similar trends were observed in flies with intact olfactory systems (Figure S5), indicating that the preference is not dependent on antennal ablation.

To control for the potential confound that the mobile ball could drift into less favorable regions of the arena – such as near walls, where flies might be less inclined to interact, we performed a positional control experiment. We fixed two immobile balls in the arena: one in the standard location used previously (ball A), and a second positioned at various distances along the axis between ball A and the arena wall (Figure S6). Interaction scores did not differ significantly across these conditions (Figure S6), indicating that wall proximity alone does not account for object preference.

### Role of hΔ neurons in the central complex in mediating object preference

Having established that flies interact differently with mobile and immobile balls, we next asked: how do flies distinguish between two objects that are visually and olfactorily identical? One key distinction is spatial position of the objects. Given that *Drosophila* are capable of forming spatial memories (Corfas, Sharma, and Dickinson 2019; Kim and Dickinson 2017; Ofstad, Zuker, and Reiser 2011), we hypothesized that flies may associate spatial locations with object properties, and that these associations could guide future interactions. To test whether spatial navigation circuits contribute to object interaction behavior, we conducted a targeted silencing screen using Kir2.1 to inhibit activity in a group of fan-shaped body neurons known as hΔ neurons. While the hΔB subtype has been shown to encode the fly’s allocentric travel direction and participate in vector-based navigation (Hulse et al. 2021; Lyu, Abbott, and Maimon 2022; Wolff et al. 2025), the roles of the remaining 13 hΔ subtypes are not well understood, and none have been directly implicated in object-related behaviors. These neurons share similar anatomical locations and projection motifs, raising the possibility that they may encode or store other spatial variables relevant to object interaction. To broadly assess the contribution of spatial computation circuits, we also silenced EPG neurons, which encode body-centered heading direction (Seelig and Jayaraman 2015) and provide input to the fan-shaped body (Lyu, Abbott, and Maimon 2022).

We screened 13 hΔ neuron subtypes, using our two-ball pinwheel-guided assay to evaluate whether silencing any subtype impaired the fly’s ability to develop a preference for the immobile ball (Figure 6A). Silencing five hΔ lines – hΔA, hΔJ, hΔM, hΔK, and hΔI – significantly reduced preference, as measured by the difference in interaction scores between the immobile (ball A) and mobile (ball B) objects (Figure 6B).

**Figure 6.**
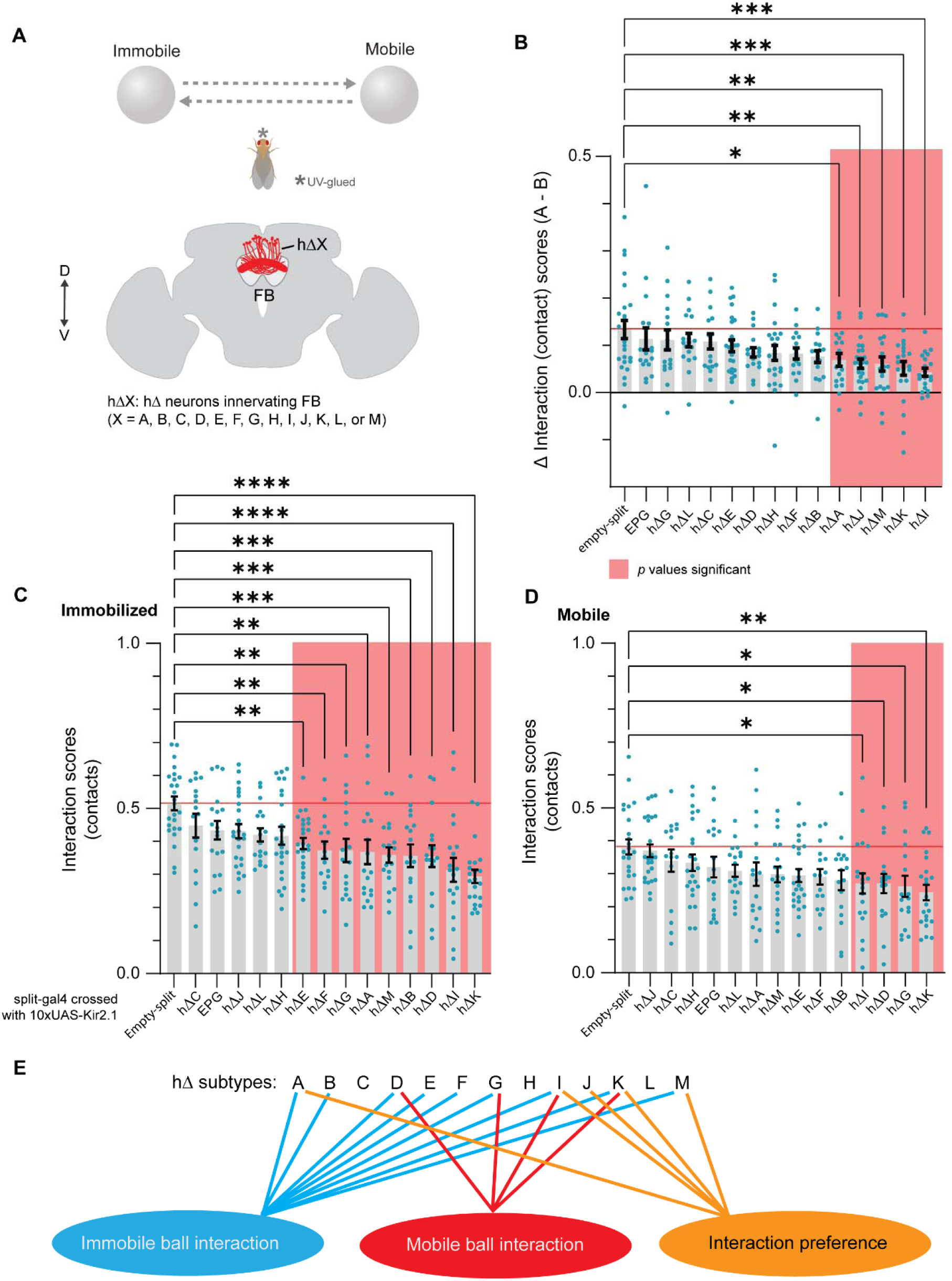
A subset of hΔ neuron subtypes influence fly-ball interaction probabilities and preference. **(A)** To identify neural substrates of object interaction behavior, we silenced 13 distinct hΔ neuron types and EPG neurons using 10xUAS-Kir2.1 and tested the resulting flies in a two-ball interaction assay. In this assay, flies were alternately guided to interact with an immobile and a mobile ball during the same session. Antennae were UV-glued to eliminate olfactory input. Each hΔ split-GAL4 line (hΔA–hΔM), as well as the EPG line (SS00090), were crossed to 10xUAS-Kir2.1. *n* values for each genotype were as follows: hΔA (18), hΔB (17), hΔC (16), hΔD (17), hΔE (26), hΔF (16), hΔG (17), hΔH (24), hΔI (20), hΔJ (25), hΔK (22), hΔL (18), hΔM (19), EPG (18). An empty-split-GAL4 × 10xUAS-Kir2.1 line was used as the control group (*n* = 25). **(B)** Silencing of five hΔ subtypes (hΔA, hΔJ, hΔM, hΔK, hΔI) significantly reduced the preference for immobile over mobile balls, as measured by the Δ interaction score (immobile – mobile). *One-way ANOVA, Bonferroni’s multiple comparisons test* was used. Significant groups are shaded in light red (*p* < 0.05). **(C)** Interaction scores for the immobile ball condition. Nine hΔ lines (hΔE, hΔF, hΔG, hΔA, hΔM, hΔB, hΔD, hΔI, hΔK) showed significantly reduced scores compared to controls (*One-way ANOVA, Bonferroni’s multiple comparisons test*). **(D)** Interaction scores for the mobile ball condition. Four groups (hΔI, hΔD, hΔG, hΔK) showed significantly reduced scores compared to controls (*One-way ANOVA, Bonferroni’s multiple comparisons test*). **(E)** Summary diagram highlighting the hΔ subtypes implicated in immobile ball interactions, mobile ball interactions, and interaction preference.

To further dissect this effect, we examined the likelihood of interaction with each ball type. Silencing hΔE, hΔF, hΔG, hΔA, hΔM, hΔB, hΔD, hΔI, or hΔK significantly reduced interactions with the immobile ball (Figure 6C, Figure S8C). In contrast, only a subset of these lines – hΔI, hΔD, hΔG, and hΔK – showed significantly reduced interaction with the mobile ball (Figure 6D, Figure S8D). These results suggest that a broader network of hΔ neurons (Figure S7A) supports interaction with immobile objects, and that these neurons may contribute to the spatial representations that underlie object evaluation. Interestingly, silencing EPG neurons had no effect on the flies’ ability to develop a preference for immobile objects, suggesting that the egocentric heading direction system is not involved in this behavior.

### hΔ neurons influence specific fly-ball interaction motifs and guidance behavior

Beyond their role in object preference, hΔ neurons also contributed to the expression of specific interaction motifs. Silencing hΔM significantly reduced the likelihood that flies would perform ball-pull maneuvers (Figure 7A). In contrast, silencing hΔJ led to an increase in ball-walk behavior, in which flies remained on the ball while walking along its surface (Figure 7B). Silencing hΔC resulted in a higher probability of jumping from mobile balls (Figure 7C, Figure S8E), though no hΔ lines significantly altered jump probability from immobile balls (Figure 7D).

**Figure 7.**
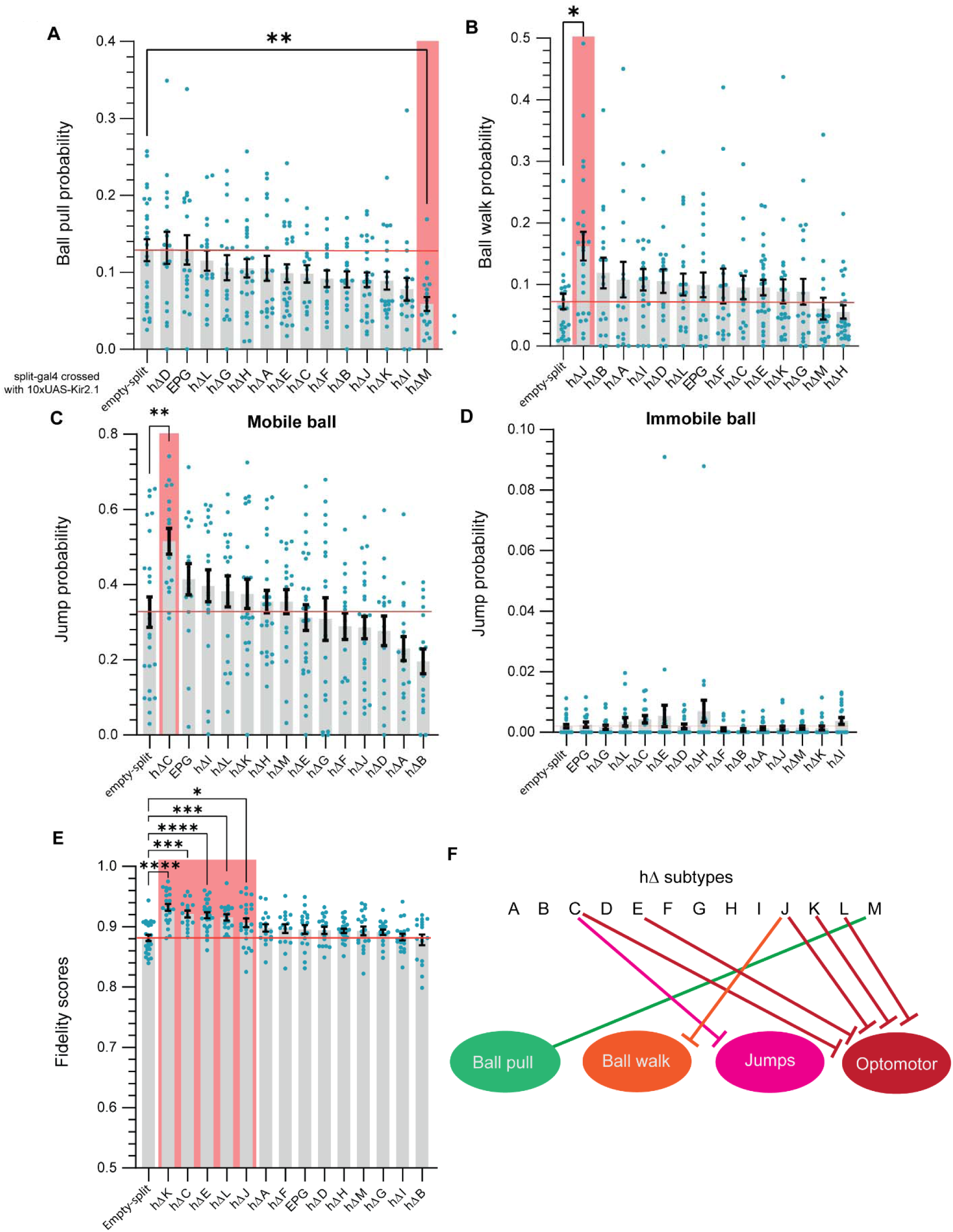
Distinct hΔ neuron subtypes modulate specific behaviors expressed during fly-ball interactions. **(A)** Ball-pull probability, defined as the proportion of contact events followed by a ball pull, across the 15 genetic conditions tested in the two-ball assay (13 hΔ subtypes, EPG neurons, and control). Silencing hΔM resulted in a significantly lower pull probability than control. *One-way ANOVA with Bonferroni’s multiple comparisons test.* Sample sizes (*n*) for each condition match those listed in Figure 6A and are identical across all panels in this figure. **(B)** Ball-walk probability, defined as the proportion of contacts resulting in a ball walk. Silencing hΔJ significantly increased ball-walk probability compared to control. *Kruskal–Wallis test with Dunn’s multiple comparisons* (control distribution was non-normal). **(C)** Jump probability for immobile balls, calculated as the number of jumps divided by the number of contacts. Silencing hΔC significantly increased jump probability. *One-way ANOVA with Bonferroni’s multiple comparisons test*. **(D)** Jump probability for immobile balls. No genotype showed a statistically significant difference from control. *Kruskal–Wallis test with Dunn’s multiple comparisons test*. **(E)** Fidelity scores, calculated as the proportion of successful visual guidance turns. Silencing hΔK, hΔC, hΔE, hΔL, hΔJ significantly increased fidelity relative to control. *One-way ANOVA with Bonferroni’s multiple comparisons test.* One outstanding outlier was removed from hΔE by ROUT method (Q = 0.1%) by GraphPad Prism. **(F)** Summary diagram indicating which hΔ neuron subtypes are required for specific motor outcomes. T-bar lines indicate inferred inhibition of behavior.

Interestingly, we also observed that silencing several hΔ subtypes – hΔK, hΔC, hΔE, hΔL, and hΔJ – led to significantly increased fidelity in following the opto-motor based visual guidance cue (Iwasaki et al. 2025). This suggests that activity in these neurons may normally compete with opto-motor responses, and that their inhibition allows flies to more readily follow externally driven visual guidance.

## Discussion

Our results demonstrate that repeated visual guidance of *Drosophila melanogaster* toward a small object promotes voluntary and sustained interaction. Flies engaged in a range of behavioral motifs – from mounting and dismounting to more complex patterns like “ball walking,” where they maintained balance while locomoting atop a moving surface. These interactions were self-initiated, indicating agency, and evolved over time, suggesting that flies progressively refine their actions through sensorimotor integration.

Unguided flies rarely interacted with the ball, consistent with prior studies showing object aversion during flight or ambiguous object engagement during walking (Maimon, Straw, and Dickinson 2008; Wehner 1972; Bülthoff, Götz, and Herre 1982). This indicates that proximity alone is insufficient to drive consistent engagement. Our results suggest that visual guidance may act as a behavioral priming mechanism – either by increasing arousal, redirecting attention, or creating a closed-loop contingency where flies gain feedback from influencing object movement. Control experiments ruled out passive visual stimulation or spatial closeness as sufficient drivers of consistent interaction, highlighting the importance of active engagement. Repeated safe exposure may also reshape object valence, reducing avoidance over time in a process reminiscent of exposure-based learning (Craske et al. 2014; Cheng, Colbath, and Frye 2019).

Among the most striking behaviors was ball walking, which emerged progressively across trials. This behavior requires balancing on a curved, unstable surface while moving – an act that likely demands the integration of mechanosensory and visual input. Its emergence over time is consistent with gradual skill acquisition, a hallmark of learning in other insect systems (Kriete and Hollis 2022; Loukola et al. 2017; Dukas 2008; Behmer 2005). One possibility is that ball walking may lie on a behavioral continuum with ball pulling, in which flies displace the ball while dismounting. Ball pulls may represent an earlier stage of motor coordination, with flies initially shifting the ball upon exit before learning to stabilize and actively locomote across it. This sequence hints at progressive refinement in motor control. Alternatively, ball walks may constitute a distinct behavioral mode, potentially reflecting object exploration, engagement, or even play-like activity. This interpretation is supported by our findings in Figure 7A,B, where silencing hΔM specifically disrupted ball pulls, whereas silencing hΔJ selectively increased ball walks without affecting ball pulls. Future experiments using circuit manipulations and high spatiotemporal tracking of limb kinematics could reveal this motor progression in much more detail, for example by testing whether flies exhibit changes in posture, gait, or limb coordination across repeated ball walks. Such data could help distinguish between reflexive displacement and intentional ball locomotion, and reveal whether flies show active balancing strategies, adaptive control, or progressive fine-tuning of movement parameters over time, hallmarks of sensorimotor learning seen in other insects (Loukola et al. 2017).

While ball walking may reflect growing motor competence, its spontaneous and persistent nature across trials also raises the intriguing possibility that it resembles play. Play-like behaviors, once thought exclusive to mammals and birds (Diamond and Bond 2003; Kaplan 2024), have been proposed in invertebrates (Burghardt 2005; Triphan, Ferreira, and Huetteroth 2025; Galpayage Dona et al. 2022). Play is generally considered to promote cognitive flexibility, sensorimotor coordination, and exploratory problem-solving. The expression of non-goal-directed, physically demanding behaviors such as ball walking may reflect similar functions in flies, even in the absence of external reinforcement.

Flies may also interpret or respond to object interactions with motivational states shaped by internal drives. Some ball engagement, particularly mounting, pushing, or “wrestling”, could recruit motor programs that overlap with courtship or mating behaviors (Yamamoto and Koganezawa 2013). Interacting with a persistently present object in a confined environment may activate latent affiliative or reproductive motor patterns, especially in socially isolated males.

Interaction patterns were also strongly modulated by object mobility. Flies exhibited a robust preference for immobile balls over mobile ones, despite being visually and olfactorily indistinguishable. This suggests that flies can detect and evaluate mechanical properties through direct physical interaction. Mechanosensory input – from structures such as campaniform sensilla or vibration-sensitive neurons (Tuthill and Wilson 2016; Lee et al. 2025), likely plays a central role. Alternatively, flies may infer stability through mismatches between expected and observed visual flow, potentially engaging circuits within the central complex to detect these discrepancies (Turner-Evans et al. 2017; Lyu, Abbott, and Maimon 2022). While this study focused on spherical objects, it remains an open question whether flies use similar strategies to evaluate and distinguish among a broader range of object types, including those with different shapes, textures, or compliance, a topic we consider promising for future investigation.

One possibility is that object stability facilitates memory formation. Spatial learning in flies is well established (Ofstad, Zuker, and Reiser 2011; Kim and Dickinson 2017; Behbahani et al. 2021), and our results suggest that learning about object features may be embedded within spatial representations. The mushroom body and fan-shaped body, which mediate associative learning and spatial integration respectively (Aso et al. 2014; Mussells Pires et al. 2024), are well positioned to support this function. Recent connectomic evidence linking mushroom body output neurons to central complex circuitry (Li et al. 2020) further supports this possibility.

From an ethological perspective, object exploration may allow flies to evaluate environmental affordances such as shelter, elevation, or navigational cues – a strategy widely seen across animals including seed-caching birds, octopuses, and rodents (Mather and Anderson 1999; Brodin 2005; Diamond et al. 2008). Flies exhibit a strong preference for elevated surfaces (Robie, Straw, and Dickinson 2010) and can orient along gravity gradients (Kamikouchi et al. 2009; Armstrong et al. 2006; Kladt and Reiser 2023), suggesting that elevation-seeking is an adaptive strategy. Interacting with a small, raised object may thus align with conserved drives for environmental monitoring and predator avoidance.

In addition, we observe that flies do not make a binary choice to engage exclusively with the immobile ball while avoiding the mobile one. Instead, their behavior suggests a relative preference shaped by cost-benefit evaluation: they tend to interact more with the object offering more favorable physical properties, while still periodically exploring the alternative. This ongoing sampling implies that the flies weigh effort or stability against novelty or uncertainty, rather than making a rigid commitment to one object type.

Mechanistically, we found that specific hΔ neurons in the fan-shaped body contribute to the preference behavior based on object mobility. Neurons most critical for object preference (hΔJ, hΔI, hΔA, hΔK, hΔM) were more interconnected than others (Figure S7A), suggesting a functionally coherent subnetwork. These neurons also received strong input from hΔB neurons, which encode allocentric travel direction (Lyu, Abbott, and Maimon 2022), implicating vector-based computation in object evaluation. Under this framework, flies may assess object mobility by comparing their own movement vector with that of the object – potentially storing the stable location of the immobile ball across trials. Objects that remain in the same position may thus afford stronger spatial memory encoding, consistent with an “object permanence” heuristic (Baillargeon 1987; Pepperberg, Willner, and Gravitz 1997). Alternatively, disrupting hΔ neuron function can alter fly’s confidence in spatial predictions (Kutschireiter et al. 2023) or motor planning, and thereby shift how flies evaluate and interact with objects. Surprisingly, silencing EPG neurons did not significantly affect object preference formation. This may reflect redundancy in the fly’s navigational system, where other sources of directional or spatial information, such as proprioceptive cues from self-motion, are sufficient to support the formation of object preference.

Beyond preference, hΔ neurons also influenced behavioral motifs. Silencing hΔJ increased ball-walking behavior. Notably, hΔJ provides substantial input to FC2 neurons (Figure S7C,D), which have been implicated in spatial goal representation (Mussells Pires et al. 2024). This connectivity suggests that hΔJ may contribute to internal goal-setting processes, and that its silencing may bias flies toward more exploratory, less goal-directed behaviors. Silencing hΔC increased jumping, which may reflect agitation or escape-like behavior (von Reyn et al. 2014; Dombrovski et al. 2023), especially given flies’ reduced willingness to contact mobile balls. Alternatively, jumping could result from low-level aggression toward an obstructive or unpredictable object. hΔC’s input from FB6A neurons, implicated in sleep regulation (Hulse et al. 2021), suggests a role in arousal or state regulation, though no changes in overall locomotion were observed (Figures S8A, S8B).

Finally, silencing specific hΔ neurons (hΔK, hΔC, hΔE, hΔL, hΔJ) enhanced flies’ ability to follow visual guidance cues. This may reflect reduced competition between internally generated goals and externally imposed cues. All but hΔK strongly influence FC2 neurons (Figure S7C,D), which encode goal heading (Mussells Pires et al. 2024). Thus, disrupting internal goal circuits may unmask latent opto-motor reflexes, improving guidance fidelity.

In summary, our findings reveal that flies use physical interaction to extract object properties, integrating visual, mechanosensory, and spatial feedback to guide future decisions. These behaviors, previously attributed largely to vertebrates, demonstrate that even small insect brains are capable of structured learning and adaptive exploration. Our experimental paradigm opens new opportunities to dissect the neuronal algorithms underlying object recognition, learning, and memory in *Drosophila*. Given the genetic tools available for circuit dissection in flies, future studies can illuminate how compact neural architectures achieve complex behavioral outcomes through the integration of sensory feedback, motor practice, and associative learning. More broadly, uncovering the principles by which flies learn about and interact with their environment may provide fundamental insights into the neural bases of exploration, memory formation, and adaptive behavior across species.

## Author contributions

Conceptualization: KI, AR

Methodology: KI, SK, CN, AR

Investigation: KI, AR

Data analysis: KI, SK, AC, CN, AR

Funding acquisition: AR

Project administration: AR

Supervision: AR

Writing – original draft: KI, AR

Writing – review & editing: KI, AR

## Resource Availability

### Lead Contact

Further information and requests for resources and reagents should be directed to and will be fulfilled by the Lead Contact, Aleksandr Rayshubskiy.

### Materials Availability

This study did not generate new unique reagents. All fly strains used are commercially available or have been previously published and are listed in the Key Resources Table.

### Data and Code Availability

All raw data and source code supporting the findings of this study have been deposited in public repositories and are freely available:

- Raw behavioral data: OSF, DOI: <u>10.17605/OSF.IO/3RDME</u>
- Analysis code repository: GitHub, https://github.com/SashaRayshubskiy/fly_ball_analysis_code

These links are also included in the Key Resources Table.

## Acknowledgements

We thank Rosy Hosking and Andrew Murray for helpful comments on the manuscript. We thank Chenxu Zhang, Chris Stokes, Kevin Gozzi, Buck Trible, Leigh Needleman, Michael Burns and Ben de Bivort for helpful discussions. We thank Rowland staff for their assistance with instrumentation. This work was funded by Rowland Institute at Harvard.

## STAR METHODS

Detailed methods are included in the online version of the paper as follows:

- **KEY RESOURCES TABLE**
- **EXPERIMENTAL MODEL AND STUDY PARTICIPANT DETAILS**

○ Fly strains & Maintenance
- **METHOD DETAILS**

○ Fly-ball interaction experimental apparatus
○ Ball immobilization
○ Two-ball assay
○ Control for potential contamination from fly droppings
- **QUANTIFICATION AND STATISTICAL ANALYSIS**

○ Fly & ball trajectory analysis
○ Ball displacement analysis
○ Interaction Score Analysis
○ Cluster Analysis of fly-ball interactive events
○ Analysis of ball walks and ball pulls (Cluster-dependent analysis)
○ Analysis of ball walks vs. ball pulls (Cluster-independent analysis)
○ Interactive jump analysis
○ “Contact” analysis
○ Halting analysis
○ Tracking artifact filtering

### Declaration of generative AI and AI-assisted technologies in the writing process

During the preparation of this work the authors used ChatGPT4.0 in order to improve language and readability. After using this tool, the authors reviewed and edited the content as needed and take full responsibility for the content of the publication.

## Supplemental Information

**Figure S1.**
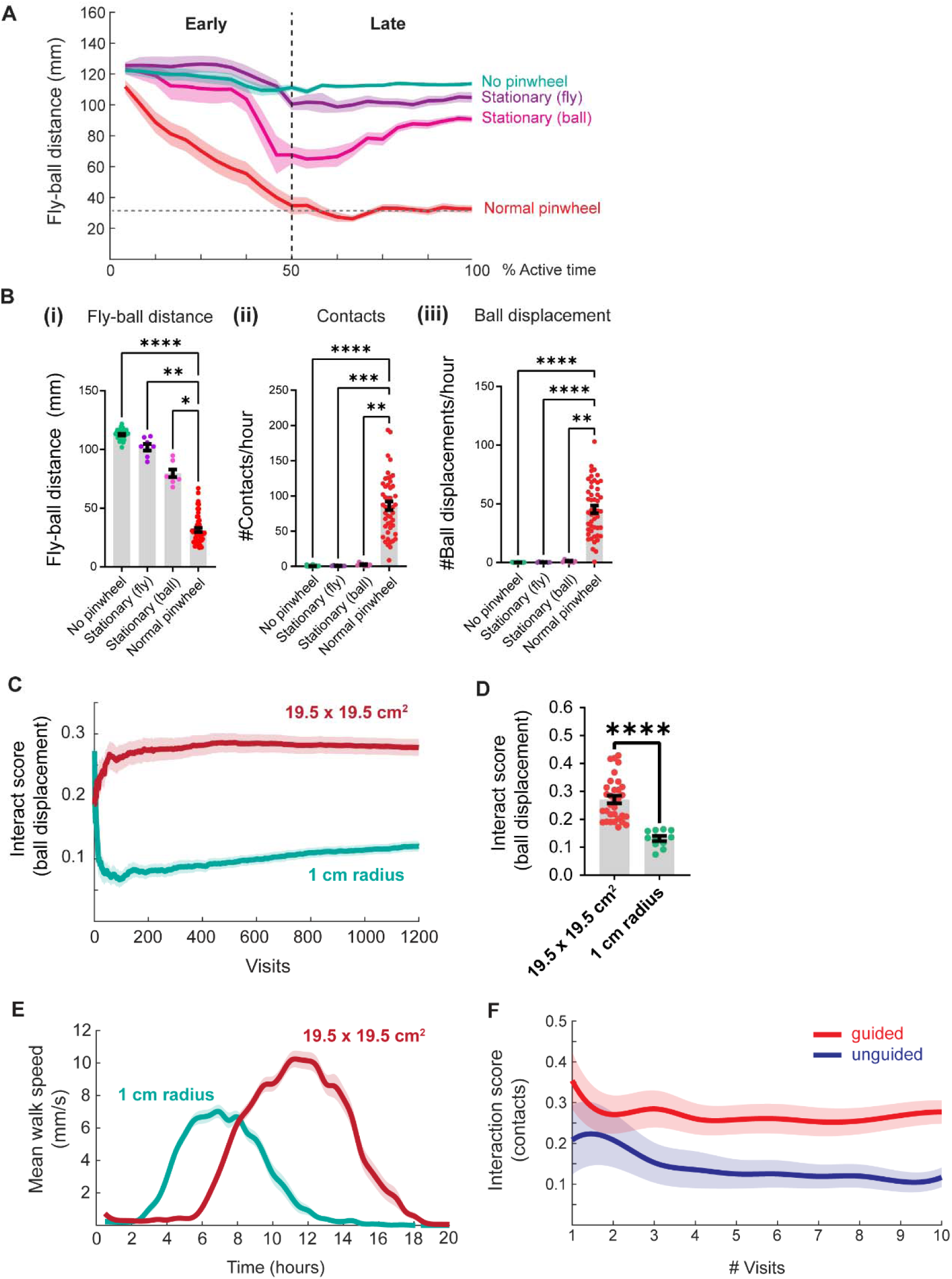
Stationary pinwheel cues do not enhance fly-ball interactions and spatial confinement alone does not explain high interaction scores observed with pinwheel guidance. To assess whether visual feedback from the rotating pinwheel influences fly-ball interactions, we compared behavior across conditions using stationary pinwheels affixed either to the fly or the ball. **A)** Fly-to-ball distance (mm) across four conditions: no pinwheel, stationary pinwheel fixed to the fly, stationary pinwheel fixed to the ball, and a dynamic (rotating) pinwheel coupled to the guided fly’s movement. Flies exposed to stationary pinwheels remained significantly farther from the balls than flies experiencing normal visual feedback (rotating pinwheel). *n* = 36 (no pinwheel), 8 (stationary-fly), 8 (stationary-ball), 48 (rotating). The final 20% of the session duration was excluded from analysis. Data are binned into 30-minute intervals. **(B)** (i) Flies in all stationary or no-pinwheel conditions maintained greater average distances from the ball compared to controls. (ii) Contact behavior was significantly reduced in all three experimental conditions relative to the rotating pinwheel control. (iii) Ball-displacement frequency was likewise reduced in all experimental groups. *Kruskal-Wallis test with Dunn’s multiple comparisons test*. *n* = 36 (no pinwheel), 8 (stationary-fly), 8 (stationary-ball), 48 (rotating). **(C)** To determine whether spatial confinement alone increases fly-ball interactions, we compared behavioral metrics across arenas of different sizes. Ball-displacement scores for the first 1,200 visits in the standard arena versus the 1-cm radius arena (*n* = 32, 11 flies). **(D)** Flies in the standard arena continued to show significantly higher interaction levels on average than those in the smaller arena. *Mann-Whitney test (two-tailed)* (*n* = 32, 11 flies). **(E)** Average walking speed of flies over time in the 1 cm radius arena versus the standard-size arena. Flies in the smaller arena become active significantly earlier, suggesting that the prolonged initial inactivity in the larger arena may reflect hesitation or environmental uncertainty associated with being in a more expansive, featureless space. Experiments were typically started in the afternoon (between 2:00 pm and 5:00 pm), meaning that the rise in locomotion speed generally occurred in the evening, approximately between 8:00 pm and 11:00 pm. The 1 cm arena curve was shifted 67 minutes to the right to align its average starting time with that of flies in the standard-size arenas. This adjustment was made to ensure comparability between experimental conditions with respect to the timing of data collection. **(F)** Contact interaction scores for guided versus unguided flies. Unguided flies made fewer visits to the ball, and during those visits, they were less likely to engage with the object.

**Figure S2:**
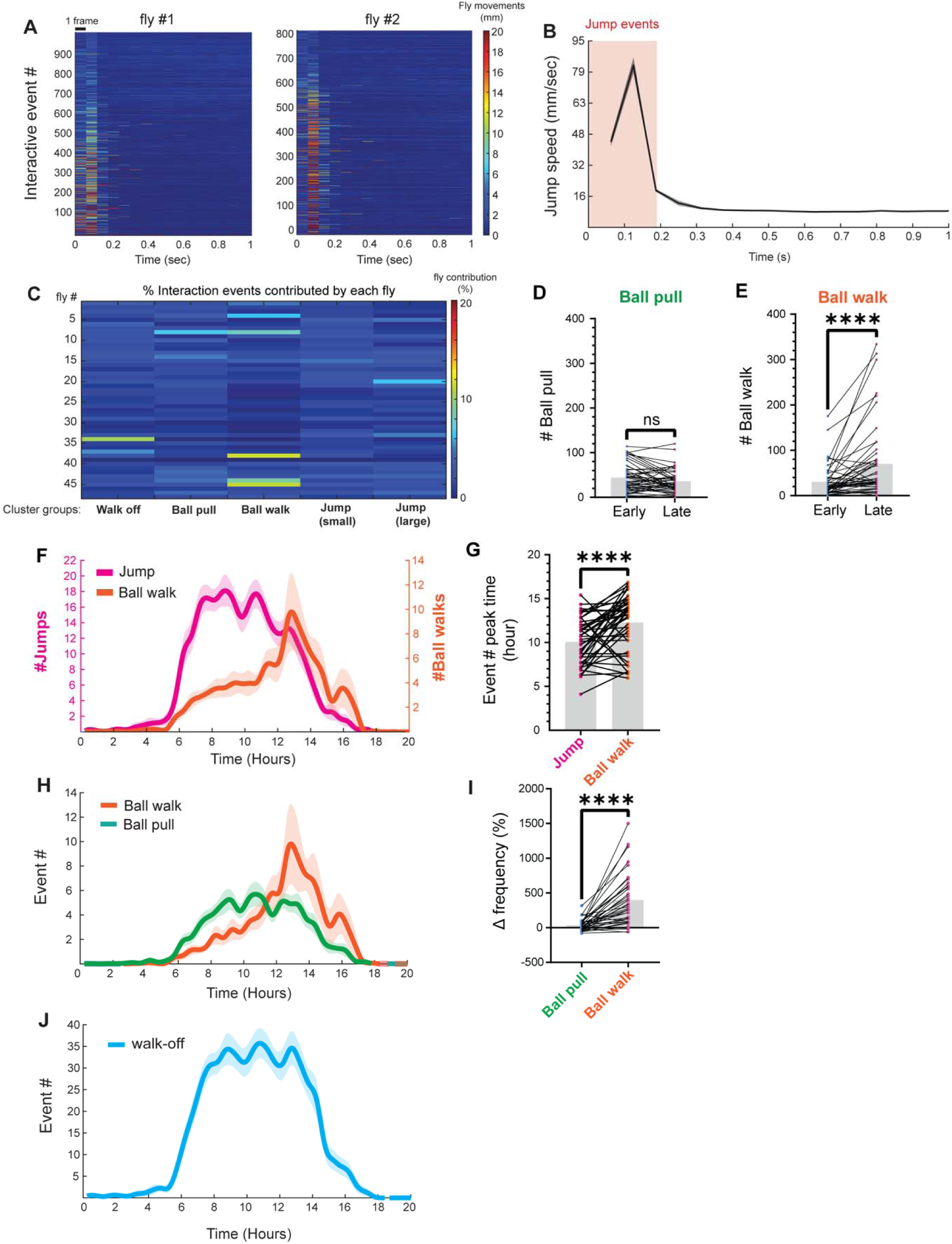
Jump-like events complete rapidly after fly-ball interactions and are similar across individuals. **(A)** Representative examples of “jump” events from two flies (#1 and #2). In both cases, the rapid movement concludes within the first three recorded frames (∼188 ms) following the fly’s exit from the ball. Note: the decrease in jump distance after approximately 600 interactions correspond to ∼9 hours of experimental time, which corresponds to a peak in overall locomotor activity (Fig S1E). We interpret this drop as habituation rather then a circadian effect. **(B)** Timing of peak jump velocity following interaction termination. Most jump events reach their maximum speed within 200 milliseconds of fly-ball disengagement (*n* = 48 flies). Number of jump events per fly = 625 ± 310 (standard deviation). **(C)** Distribution of event contributions across flies. Individual flies contributed comparably to each of the five behavioral motif clusters described in Figure 2, with no fly accounting for more than 12% of the events in any single cluster. *Cluster-dependent analysis supports differential temporal dynamics between ball-pull and ball-walk interactions*. **(D)** Frequency of ball-pull events does not differ significantly between the early and late phases of the active period (*Wilcoxon matched-pairs signed-rank test*, two-tailed; *n* = 48). **(E)** In contrast, ball-walk frequency increases significantly in the second half of the experiment (*Wilcoxon matched-pairs signed-rank test*, two-tailed; *n* = 48). **(F)** Time-course comparison of jump and ball-walk frequencies. Ball-walk events gradually rise and peak sharply around 12 hours, while jump events peak earlier and decline gradually thereafter. Jumps are defined as fly displacements >5 mm (≥2 body lengths) and ≤75 mm following dismount from a ball. Jumps defined here includes both “small” and “large” jumps defined in Figure 2. Data are binned in 30-minute intervals (*n* = 48 flies). **(G)** On average, ball-walk frequency peaks significantly later than jump frequency (*Wilcoxon matched-pairs signed-rank test*, two-tailed; *n* = 47 flies; one fly excluded due to absence of ball-walk events). **(H)** Cluster-dependent analysis confirms the overall pattern observed in Figure 3B: ball-walk frequency increases over time more than ball-pull frequency (*n* = 48; 30-minute bins). The analysis of temporal developments described in Figure 3B was re-conducted using event numbers quantified by the cluster analysis (Figure 2). **(I)** The temporal increase in ball-walk frequency is significantly greater than that of ball-pull frequency. Outlier removal (ROUT method, Q = 0.1%) excluded 12 flies from analysis for clarity (*n* = 36). The result remains significant with or without outliers (*p* < 0.0001, ****; *Wilcoxon matched-pairs signed-rank test*, two-tailed). **(J)** Time course of ‘walk off’ events associated with cluster analysis in Figure 2.

**Figure S3.**
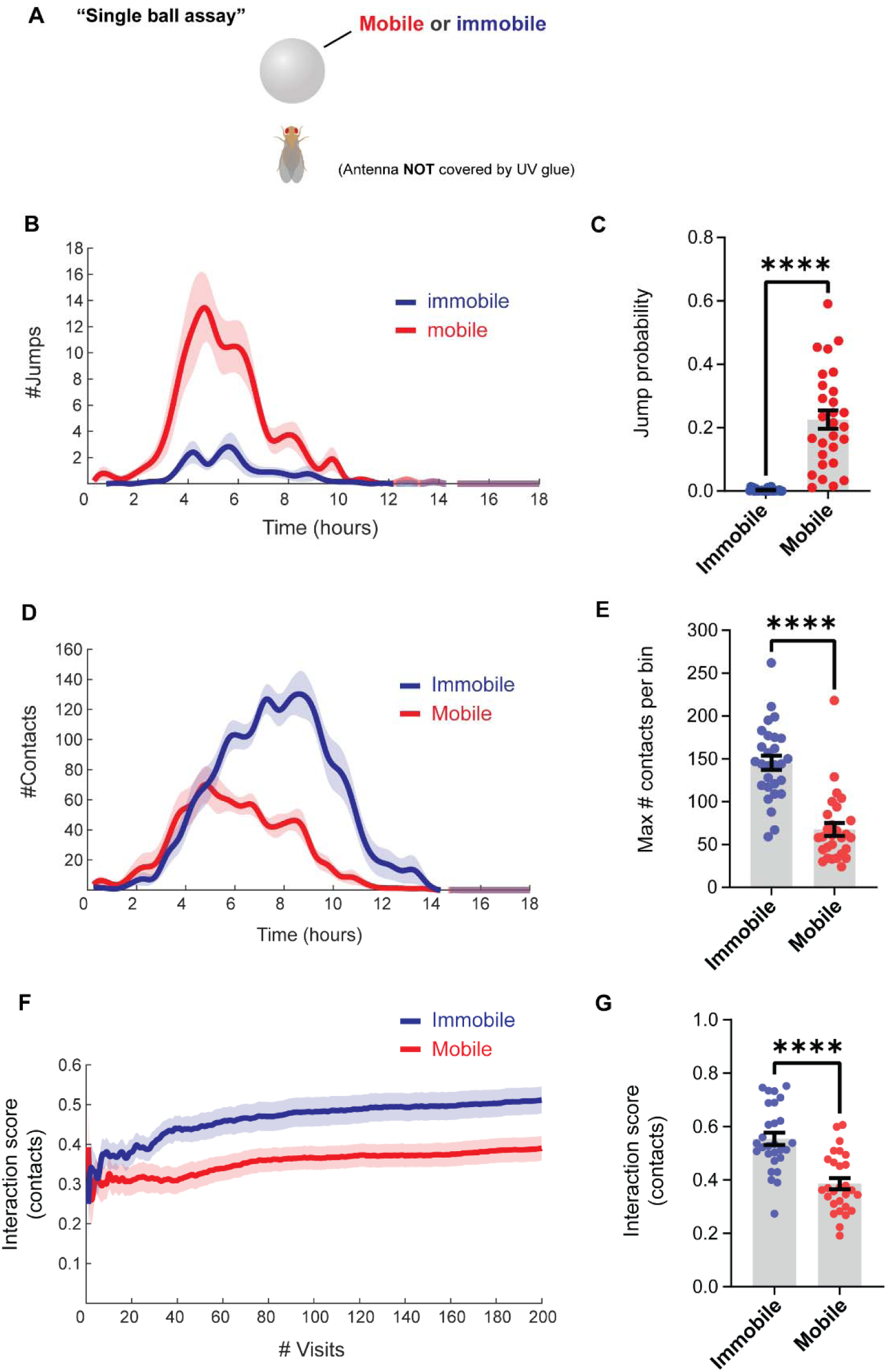
Flies exhibit a preference for immobilized balls also in the presence of intact olfactory input. This control experiment (without antennal UV-gluing) tested whether flies exhibit the same mobility-dependent interaction bias when olfactory input is intact. In contrast, flies in Figure 4 had their antennae glued to eliminate the possibility of distinguishing between the two balls based on olfactory or other antennal cues. **(A)** Schematic of the single-ball assay: flies were guided to interact with either a mobile or glued (immobile) ball. Unlike in Figure 4, flies in this experiment retained normal antennal (olfactory) function. **(B)** Time course of jump frequency for mobile versus immobile ball conditions (30-minute bins; *n* = 28 flies per group). **(C)** The probability of jumping after interaction is significantly higher for mobile balls (*Mann– Whitney test*, two-tailed; *n* = 28 flies per group). **(D)** Contact frequency over time for each condition (*n* = 28 flies per group). **(E)** The maximum number of contacts per 30-minute bin is significantly higher for immobile balls (*Mann–Whitney test*, two-tailed, *n* = 28 flies per group). **(F)** Comparison of interaction scores (contacts per visit) for mobile and immobile ball conditions (*n* = 28 flies per group). **(G)** Final interaction scores are significantly higher when flies interacted with immobilized balls, indicating a consistent bias toward contacting immobile objects (*unpaired t-test*, two-tailed; *n* = 28 per group). Global scores across the entire session were used for analysis to avoid the use of arbitrary visit windows.

**Figure S4.**
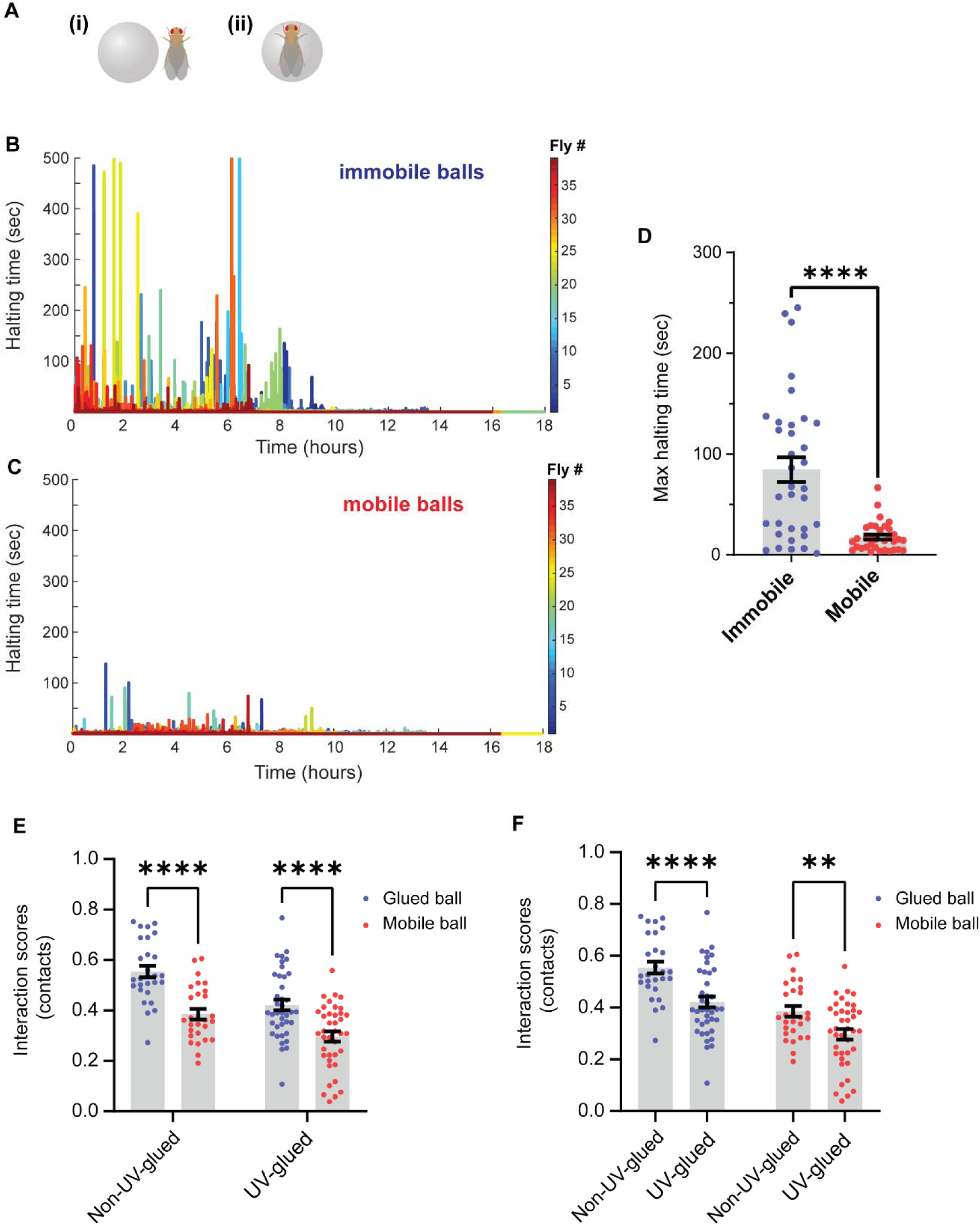
Flies spend significantly more time near immobile balls and flies preferentially interact (contact) with immobile balls regardless of antennal UV-gluing. **(A)** Representative images of flies halting either beside (i) or on (ii) balls during the experiment. **(B–C)** Total time each fly spent by or on the ball across the experiment for (B) immobilized and (C) mobile ball conditions. Each trace represents a different fly (*n* = 39 flies per group). **(D)** Maximum halting duration (time spent on or near the ball) was significantly greater for immobilized balls. *Mann–Whitney test*, two-tailed (n = 39 flies per group). Five outliers were excluded using the ROUT method (Q = 0.1%, GraphPad Prism v10.0); the *p*-value remains significant with or without outliers (****, *p* < 0.0001). This behavior could be due to a reduction of environmental uncertainty afforded by the immobile ball. **(E)** Flies exhibited significantly higher interaction scores (total # contacts/ total # visits) for immobilized balls compared to mobile balls, regardless of whether their antennae were UV-glued or left intact. Global interaction scores (total # contacts/ total # visits) across the full session were used for analysis. *Two-way ANOVA with Bonferroni’s multiple comparisons test*; *n* = 28 flies (non-UV-glued), 39 flies (UV-glued). **(F)** Overall interaction scores (contacts) for both ball conditions were significantly lower in flies with UV-glued antennae. This reduction may result from loss of olfactory input or from blocking antennal chordotonal neurons. Additional experiments are needed to distinguish between these possibilities. *Two-way ANOVA with Bonferroni’s multiple comparisons test*; *n* = 28 flies (non-UV-glued), 39 flies (UV-glued).

**Figure S5:**
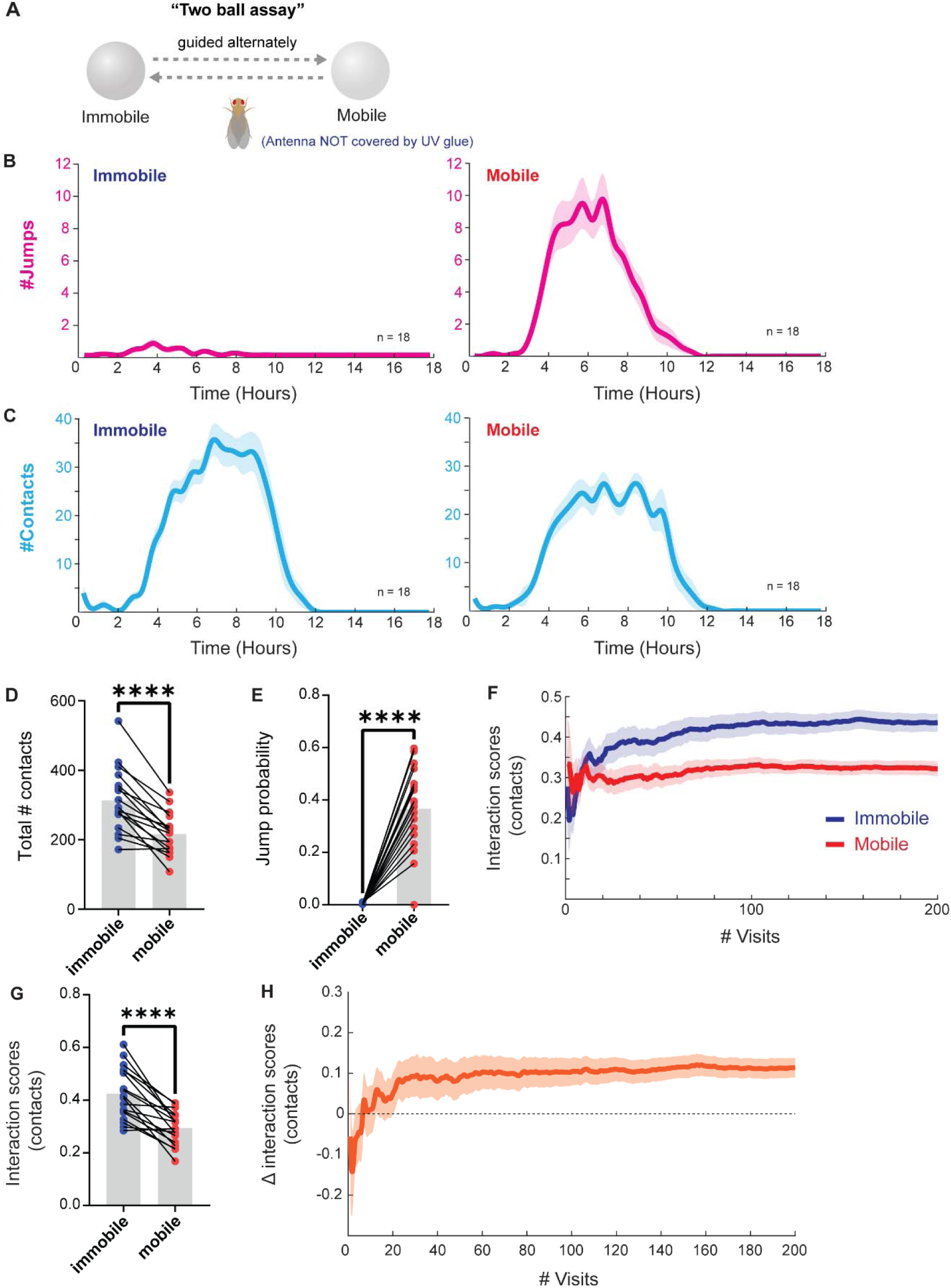
Flies develop a bias toward immobilized balls when alternately interacting with mobile and immobile balls, also with intact olfactory input. This control experiment mirrors the two-ball assay described in Figure 5, but without antennal UV-gluing. **(A)** Schematic of the two-ball assay. Test flies were alternately guided to interact with a mobile and an immobilized (glued) ball. Antennae were left intact (no UV glue). **(B)** Time course of jump frequencies for mobile versus immobile balls (*n* = 18 flies per group; 30-minute bins). The y-axis indicates the number of events per 30-min time bin. **(C)** Time course of contact frequencies for mobile versus immobile balls (*n* = 18 flies per group; 30-minute bins). The events of contacts and jumps are not mutually exclusive. Every jump occurs after an initial contact with the ball. **(D)** Total number of contacts was significantly higher for immobile balls (*paired t-test*, two-tailed; *n* = 18 flies). **(E)** The probability of jumping after interaction was significantly higher for mobile balls (*Wilcoxon matched-pairs signed-rank test*, two-tailed; *n* = 18 flies). **(F)** Interaction scores (contacts) for both conditions (*n* = 18 flies) during the first 200 visits. **(G)** Interaction scores were significantly higher for immobilized balls (*paired t-test*, two-tailed; *n* = 18 flies). **(H)** Δ interaction score (immobile – mobile) plotted as a function of number of visits. A stable preference for immobilized balls emerged after approximately 10 visits on average, following an initial dip (*n* = 18 flies).

**Figure S6.**
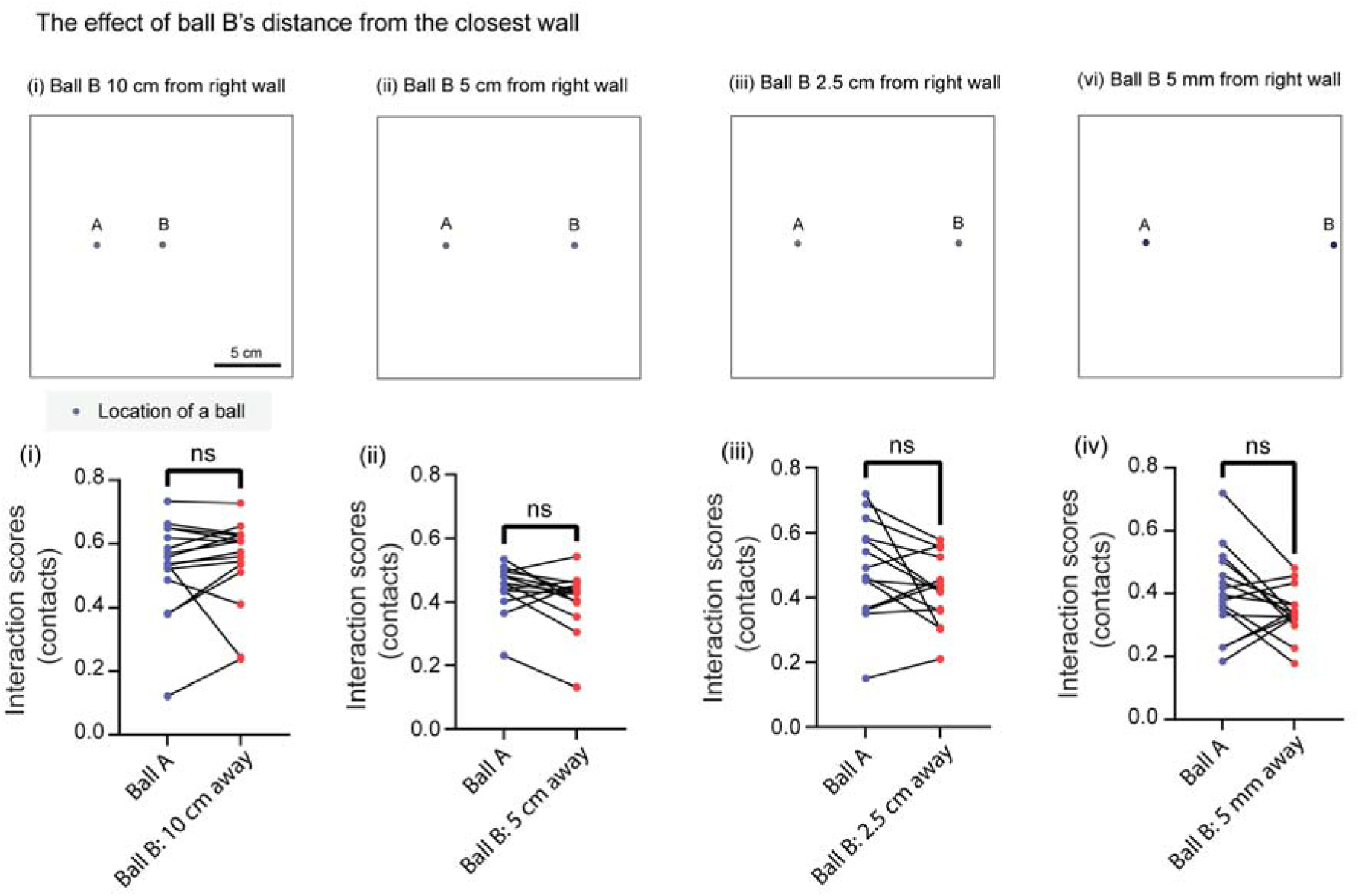
Ball location does not influence interaction scores (contacts) in two-ball experiments. To determine whether ball position in the arena affects interaction behavior, we conducted four two-ball assays in which the location of Ball B (glued) was varied across the arena (conditions i– vi), while Ball A (also glued) remained fixed. Each experiment was analyzed using a paired comparison of interaction scores (contacts per visit) for Ball A versus Ball B. No significant differences were observed in any condition. *Wilcoxon matched-pairs signed-rank tests* were used for conditions (i) and (ii), and *paired t-tests* for conditions (iii) and (vi) (*n* = 16, 14, 14, 15 respectively). The two-ball assay enabled controlled within-subject comparisons.

**Figure S7.**
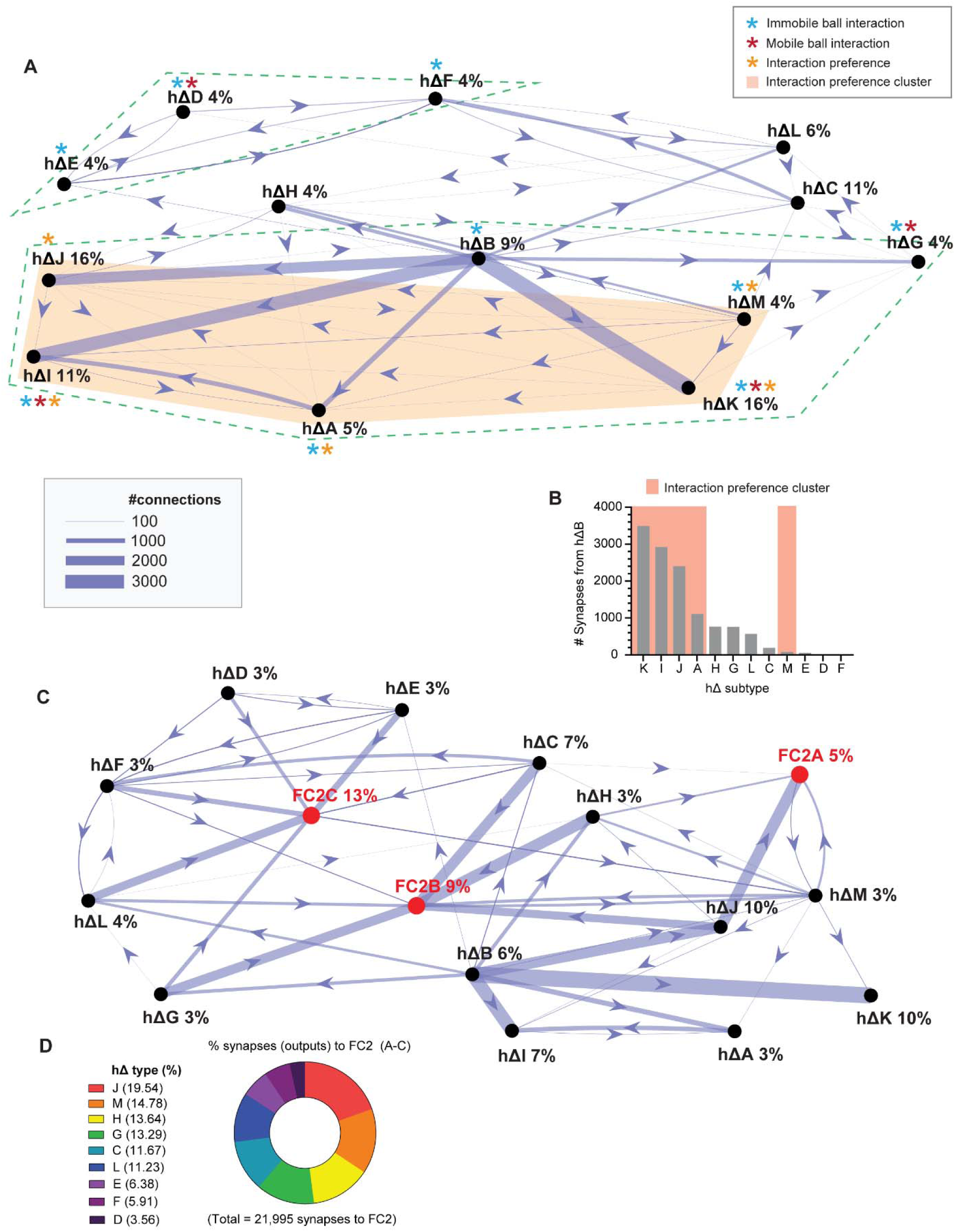
Connectivity among hΔ neuron subtypes implicated in fly-ball interaction behaviors and hΔ neuron subtypes projecting to FC2 neurons. **(A)** A connectivity map was generated using connectivity data in Codex (https://codex.flywire.ai/) (Matsliah et al. 2023; Dorkenwald et al. 2024; Schlegel et al. 2024) for all 13 hΔ neuron subtypes tested in the behavioral screen. Nodes were assigned unique identifiers, and the directed graph was generated using MATLAB’s digraph() function. Edge thickness was scaled proportionally to connection strength based on the relative weights. Neuron subtypes identified as significant for specific behavioral outcomes are color-coded: blue for immobile ball interaction, red for mobile ball interaction, and orange for interaction preference (Δ interaction score). Top 50 connections are shown with minimum five synaptic connections. The dotted green line shows a connected subgroup of hΔ neurons that play a role in ball interactions, with at least one of the asterisks. **(B)** Synaptic outputs from hΔB neurons onto downstream partners shown in the network in (A), based on synapse counts. **(C)** Network diagram showing synaptic connectivity between hΔ neuron subtypes and FC2 neurons (types A–C). Connections were visualized using Codex (FlyWire) with a ForceAtlas2/Dynamic layout. Only the top 50 connections are shown, with a minimum threshold of 5 synapses per connection. **(D)** A pie chart showing the proportions of synaptic output from hΔ neuron subtypes to FC2 neurons (types A–C) as represented in (A).

**Figure S8.**
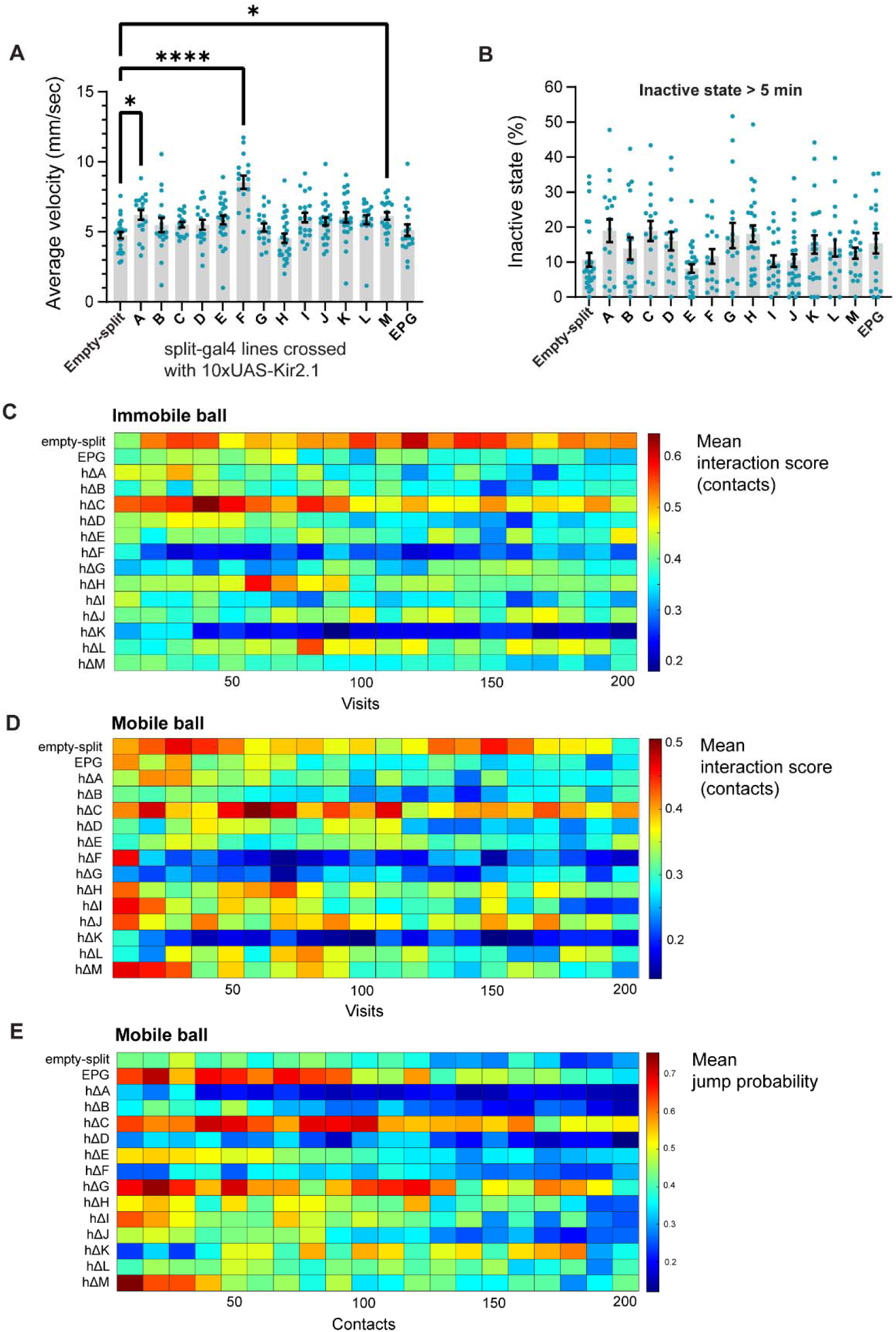
hΔ neuron manipulations do not impair general locomotion or induce prolonged inactivity. **(A)** Average locomotor velocity during active periods for each experimental group tested in the hΔ screen. No group showed reduced speed relative to the control. Three groups – hΔA, hΔF, and hΔM – exhibited significantly higher locomotor speeds than control. *One-way ANOVA with Bonferroni’s multiple comparisons test.* The sample sizes are as follows: hΔA (18), hΔB (17), hΔC (14), hΔD (19), hΔE (26), hΔF (16), hΔG (17), hΔH (24), hΔI (20), hΔJ (25), hΔK (22), hΔL (18), hΔM (19), EPG (18). An empty-split-GAL4 × 10xUAS-Kir2.1 line was used as the control group (*n* = 25). **(B)** Proportion of time spent in inactive states, calculated using two thresholds: inactivity lasting >5 minutes. No experimental group differed significantly from control. Sleep duration was defined as 5 min or more of inactivity (Shaw et al. 2000). *Kruskal–Wallis test with Dunn’s multiple comparisons test.* The sample sizes for (i) and (ii) are as follows: hΔA (18), hΔB (17), hΔC (16), hΔD (17), hΔE (26), hΔF (16), hΔG (17), hΔH (24), hΔI (20), hΔJ (25), hΔK (22), hΔL (18), hΔM (19), EPG (18). An empty-split-GAL4 × 10xUAS-Kir2.1 line was used as the control group (*n* = 25). **(C)** The mean interaction scores (contacts) of the immobile ball condition for the first 200 visits for hΔ screen as described in Figure 6 & 7. The sample size is 24, 24, 18, 17, 16, 16, 25, 16, 17, 24, 19, 25, 21, 17, 19, respectively. The bin size for the heatmap is 10 visits. **(D)** The mean interaction scores (contacts) of the mobile ball condition for the first 200 visits for hΔ screen. The sample size is 24, 24, 18, 17, 16, 16, 25, 16, 17, 24, 19, 25, 21, 17, 19. **(E)** The mean jump probability (# jump events/ # contact events) of the mobile ball condition for the hΔ screen. The sample size is 15, 5, 11, 9, 11, 5, 15, 9, 5, 9, 8, 19, 3, 10, 7.

**Video 1. An example of a ball pulling behavior, recorded from a side view**

**Video 2. Another example of a ball pulling behavior, recorded from a side view**

**Video 3. An example of a ball walking behavior, recorded from a top-down view**

**Video 4. An example of a ball walking behavior, recorded from a top-down view**

**Video 5. An example of a ball walking behavior, recorded from a side view**

**Video 6. An example of a ball walking behavior, recorded from a side view**

**Video 7. An example of a ball walking behavior, recorded from a side view**

**Video 8. An example of a jump behavior, recorded from a side view**

**Video 9. An example of a walk-off behavior, recorded from a side view**

In these videos, the pinwheel visual guidance cue was removed during recording to prevent interference with fly and ball tracking. A blue circle in the upper left corner indicates the status of the guidance cue: it appears when the cue is active and disappears when the cue is off. Several videos were presented for the ball-walk motif to provide multiple perspectives on this highly complex fly–ball interaction motif.

## STAR⍰ METHODS

## KEY RESOURCES TABLE

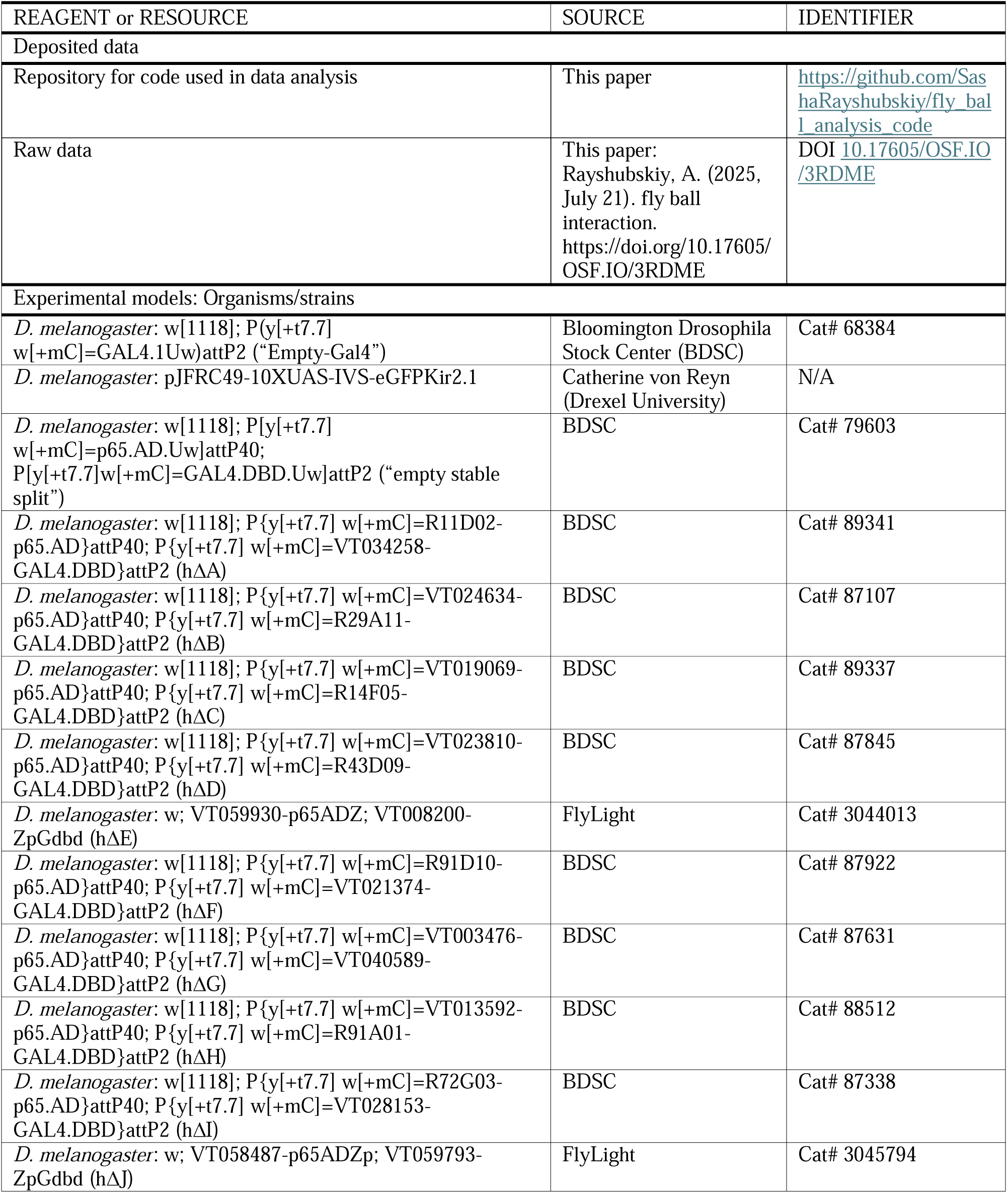
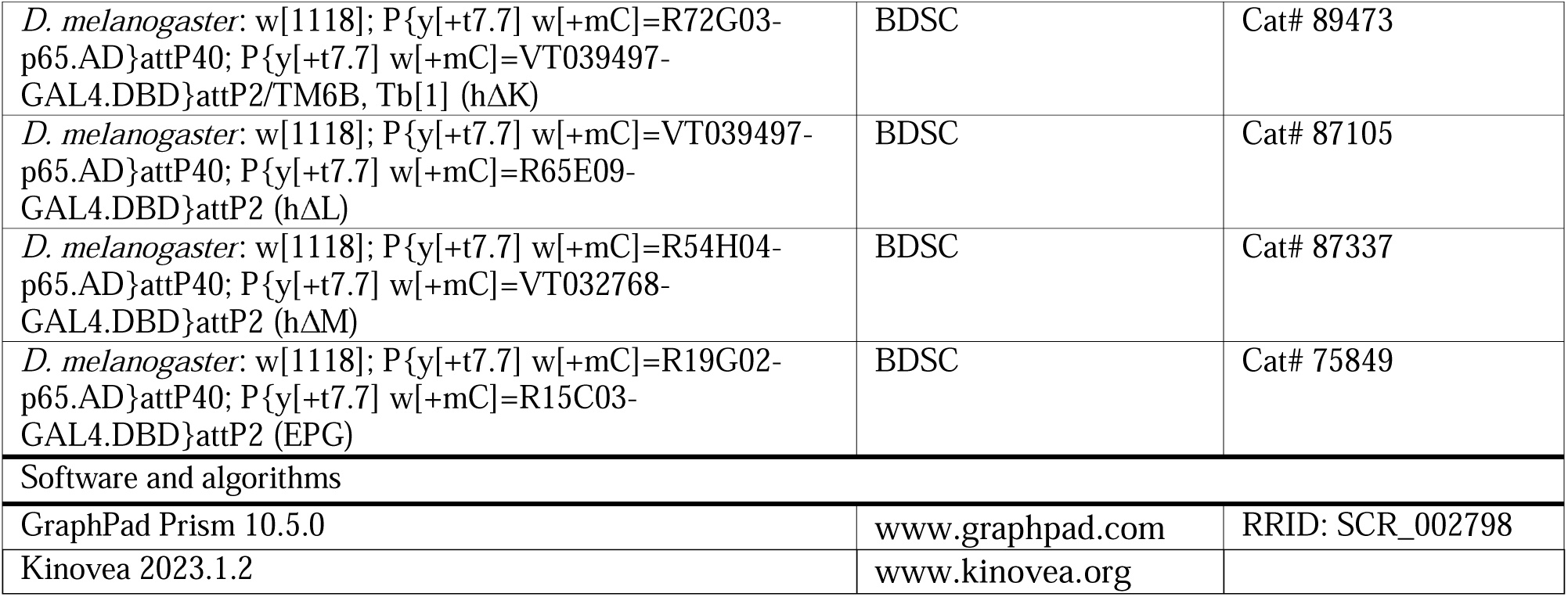

## EXPERIMENTAL MODEL AND STUDY PARTICIPANT DETAILS

### Fly strains & Maintenance

Flies were reared on standard cornmeal medium (Corn syrup–soy formulation, Archon Scientific) under a 12:12 h light:dark cycle at 25 °C. Stocks were maintained in vials within environmental chambers (Biocold Environmental Inc., Insect Environmental Chamber; Percival Scientific, Model LED-36L2) with relative humidity set to 50%. Genotypes of adult flies used in each experiment are detailed in the KEY RESOURCES TABLE. Experiments were typically started in the afternoon between 2 pm and 5 pm. Male flies were tested at one day post eclosion for all experiments. Male flies were used in the current study due to a technical reason. Male flies are smaller in body length compared to the balls used in the experiments and therefore easier to track the male flies and balls independently. Testing female flies is one of the future goals of our study.

Empty-Gal4 male flies were used for Figures 1 – 5 and Supplemental Figures S1-S6. For the hΔ neuron genetic screen (Figures 6-7 & S7-S8), pJFRC49-10XUAS-IVS-eGFPKir2.1 flies were crossed with flies from each of the thirteen hΔ split-gal4 lines, EPG split-gal4 line, and “stable empty split” line (control).

## METHOD DETAILS

### Fly-ball interaction experimental apparatus

Fly–ball interaction experiments were conducted in a custom-built behavioral chamber (43 × 43 cm), subdivided into four individual arenas (19.5 × 19.5 cm each) using 8-mm plastic dividers, as previously described (Iwasaki et al. 2025). Male flies were reared one day post-eclosion from vials containing either Formula 4-24 (Carolina Biological Supply) or Corn Syrup/Soy food (Archon Scientific). All male flies were tested at one day after eclosion. Prior to each experiment, a 2.38-mm diameter, 10-mg white Delrin acetal resin ball (McMaster-Carr #9614K51) was placed at the center of each arena. Flies were briefly anesthetized on ice (20–30 seconds) and positioned in a corner of the arena. An acrylic cover (6 mm thick) was placed atop the chamber to prevent flight, setting the vertical clearance to 5 mm. The arena walls and underside of the cover were treated with SigmaCote (Sigma-Aldrich SL2-25ML) to prevent wall climbing.

Fly behavior was recorded at 16 fps using a FLIR Blackfly S camera (BFS-U3-63S4M). Flies were visually guided toward the ball using a rotating pinwheel cue (480 nm, 2.4–2.6 μW/mm²) projected by an Optoma ZH450. The cue was automatically turned off when the fly entered a 5-mm radius “close-enough zone” around the ball and reactivated upon exit. See our previous study for pinwheel guidance parameters (Iwasaki et al. 2025). Each experiment continued until the fly died. The arena surface was leveled to within 0.05° using a dual-axis digital protractor (TLL-90S, Jingyan Instruments) to prevent unintended ball movement.

### Ball immobilization

For the immobilized ball condition, each ball was affixed to the arena floor using a small amount, approximately (0.5 – 1.0 µl) of School Glue (Elmer’s). A thin line of glue was first applied to a sheet of 3M Post-it paper, and each ball was lightly touched to the glue using fine-point tweezers (2A-SA, Romain) before being placed in the arena (Figure 4A). All glued balls were left to dry for at least 30 minutes prior to the start of the experiment. The amount of glue used was sufficient to prevent ball displacement by the fly, while keeping the vertical height nearly identical to that of a free ball (Figure 4A). If a glued ball exhibited any movement during the experiment, suggesting incomplete adhesion, the experiment was excluded from further analysis.

### Two-ball assay

In the two-ball assay, a mobile and an immobilized ball were placed 10 cm apart at the center of each test arena. Flies were guided alternately to the two balls using the visual pinwheel navigation cue. Once a fly exited the close-enough zone surrounding one ball, the guidance cue redirected it toward the other. After each experiment, each fly’s data was checked for tracking errors as follows. First, if the tracked movement of a glued ball becomes greater than 10 pixels, the fly was excluded from analysis. If the tracked movement of either ball exceeds 50 pixels per frame (198 mm/sec), the fly was also excluded from analysis. Flies were also removed from analysis if the two balls collided during the experiment and the identities of the two balls were accidentally swapped due to tracking errors. There were no other visual landmarks provided in the area to help flies discern mobile or immobilized balls.

### Control for potential contamination from fly droppings

To assess whether the observed fly-ball interaction motifs, such as ball pulling and ball walking, might be influenced by the accumulation of fly excreta on the ball surface, we conducted a post-experiment contamination control. Twenty 1-day-old male flies were allowed to interact with balls until death. At the conclusion of each experiment, the ball was examined under a stereo zoom microscope (Leica S7 E) for the presence of droppings. Only 1 of the 20 balls showed any sign of contamination: a single fecal spot approximately 0.4 mm in radius. These results indicate that accumulation of fly droppings is rare under our experimental conditions and is unlikely to account for the progressive changes in fly–ball interaction behavior observed over time. All flies displayed ball walking behavior in this experiment.

## QUANTIFICATION AND STATISTICAL ANALYSIS

### Fly & ball trajectory analysis

Fly and ball trajectories were extracted from x,y coordinates recorded during the experiment, as described in our previous study (Iwasaki et al. 2025). Behavioral data were recorded at 16 frames per second using a FLIR Blackfly S camera (BFS-U3-63S4M). Inactive flies that do not show an average locomotor speed greater than 2.22 mm/sec in any of 15-minute time bins during the experiment were excluded from analysis.

### Ball displacement analysis

Ball displacement events were identified by first detecting “entry time points” – defined as the time a fly entered the “close-enough zone” (within a 5-mm radius) around a ball. During the experiment, flies were repeatedly guided to this zone to promote fly–ball interactions. To detect ball displacement events, ball movement was assessed during the 2.5 seconds following each entry point, based on the observation that most fly-induced displacements occur within this window. A ball displacement event was defined as a displacement of more than 1 mm occurring over at least four consecutive frames (i.e., >250 ms). To minimize false positives due to tracking noise, sequences consisting only of single-pixel displacements (0.25 mm) were excluded. The mean number of ball displacement events was calculated by averaging event counts within 30-minute time bins across all test flies.

### Interaction Score Analysis

Cumulative interaction scores for contacts or ball displacements were calculated by dividing the total number of contacts or ball displacement events by the number of visits, where a “visit” was defined as an instance in which the fly entered the close-enough zone (within 5 mm of the ball). Maximum interaction scores were determined using a sliding window of 50 visits. For each fly, the highest interaction score observed within any 50-visit window was reported as the maximum.

### Cluster Analysis of fly-ball interactive events

A total of 29,982 fly-ball interaction events were analyzed from 48 *Empty-GAL4* male flies. Events were clustered into five groups using the k-means algorithm (MATLAB), based on three metrics: pre-exit ball movement, post-exit ball movement, and post-exit fly movement (Figure 2). We chose the k-means algorithm for its simple implementation and sufficient capability to capture the diversity of interactive strategies observed in our large dataset. Interactive events were defined as periods during which a fly entered and remained within a 5.0-mm radius “close-enough zone” around a ball. The start of an event was marked by the fly’s first contact with the ball; exit was defined as the moment when the fly–ball distance exceeded zero. Post-exit fly movements exceeding velocity – 532 mm/s – were excluded as tracking artifacts. Pre- and post-exit ball movements were measured in the 2 seconds before and after the fly left the ball. Displacements <0.5 mm per 63 ms frame (2 pixels) were filtered out. Post-exit fly movement was calculated over a 188 ms window, based on prior data showing that most exit-related jumps occur within this period (Figures S2A,B).

Five clusters were chosen to isolate “walk-off” events (minimal fly and ball movement) and reduce motif redundancy (Figure 2A). Each cluster was defined by distinct kinematic features and assigned a putative behavioral label (Figure 2B):

- *Walk-offs*: minimal ball and post-exit fly movement
- *Ball pulls*: large post-exit ball movement, low post-exit fly movement
- *Ball walks*: large pre-exit ball movement, low post-exit movement
- *Small jumps*: post-exit fly speed >53 mm/s
- *Large jumps*: post-exit fly speed >133 mm/s

Cluster composition was evenly distributed across flies; no individual contributed more than 12% of events to any single motif (Figure S4C). Overall distribution was as follows: walk-offs (46%), small jumps (19%), ball walks (14%), ball pulls (13%), and large jumps (8%) (Figure 2C).

### Analysis of ball walks and ball pulls (Cluster-dependent analysis)

To examine changes in behavior over time, we compared the frequency of cluster-defined ball walk and ball pull events between the first and second halves of each experiment (Figures S2H, S2I). For this analysis, only periods during which flies were actively moving (“active times”) were included. Each experiment was divided into 15-minute bins, and total fly movement was calculated per bin. Displacements <0.5 mm per frame (63 ms) were excluded to minimize tracking noise. Bins in which the average fly speed exceeded 2 mm/s were classified as “active.” The full active period was defined as the interval between the start of the first active bin and the end of the last active bin. Ball walk and ball pull event counts were then compared between the early and late halves of this active window to assess temporal shifts in behavior.

### Analysis of ball walks vs. ball pulls (Cluster-independent analysis)

Ball walks and ball pulls were also analyzed using a cluster-independent approach based on kinematic criteria (Figure 3).

*Ball walks* were defined by continuous ball displacement while the fly remained on the ball. To qualify, the total displacement had to exceed 2.5 mm, excluding movements likely to result from tracking noise or center-of-mass shifts (which can occur due to the fly’s body weight). Additionally, the fly had to remain on the ball for at least one frame (63 ms). Total displacement was calculated by summing inter-frame distances throughout the interaction.

*Ball pulls* were defined by post-exit displacement of the ball. While the fly remained on the ball, movement had to be <2.5 mm. A ball pull was then detected if displacement after the fly exited the ball exceeded 3 mm. Pull magnitude was calculated as the total distance moved between the moment of fly exit and when the ball came to rest.

To assess behavioral changes over time, active periods were split into early and late halves for comparison.

### Interactive jump analysis

Interactive jumps, defined as rapid fly movements immediately following ball interaction, were identified as events in which the fly’s speed exceeded two body lengths (5mm or equivalently 27 mm/s during the first three frames – 188 ms) after exiting the ball. To exclude tracking artifacts, fly movements exceeding 532 mm/s were filtered out. To examine how jumping behavior evolved over time, the number of jumps was averaged across all flies within each 30-minute time bin.

### “Contact” analysis

A “contact” was defined as a period during which the fly maintained any physical contact with the ball, with a fly–ball distance of zero. The event began when the fly first touched the ball and ended when it exited (i.e., the fly–ball distance became greater than zero). That means, during a “contact” event, a fly may climb the ball, remain on the ball, or simply touch the ball. This metric was analyzed to specifically measure the fly’s voluntary engagement with the ball, not its ability to surmount the ball.

### Halting analysis

Halting was defined as any period during which a fly remained stationary on top of or beside a ball during an interaction. This analysis was based on fly–ball distance measurements to identify when and where a contact occurred. The total duration of each interaction in which the fly remained in contact with or immediately adjacent to the ball was recorded as halting time.

### Tracking artifact filtering

Flies with tracking errors in which fly-ball distances abruptly dropped to zero despite the flies being nowhere near the balls prior to the “false contacts” were excluded from the analysis. Flies tested in the two-ball assay with tracking errors in which the identities of the two balls were accidentally switched were also excluded from further analysis. To reduce the impact of tracking errors, data from all analyses were filtered as follows: abrupt drops in fly–ball distance, fly trajectory, or ball trajectory greater than 10 mm within a single frame (63 ms) were removed. Adjacent bins containing similar high-magnitude movements over ≥10 frames (625 ms) were also excluded to account for surrounding tracking instability. Data exclusion criteria specific to individual analyses are described in the corresponding sections above.

## References

Alekseyenko, Olga V, Yick-Bun Chan, Maria de la Paz Fernandez, Torsten Bülow, Michael J. Pankratz, and Edward A Kravitz. 2014. ‘Single Serotonergic Neurons that Modulate Aggression in Drosophila’, Current Biology, 24: 2700–07.

Armstrong, J. D., M. J. Texada, R. Munjaal, D. A. Baker, and K. M. Beckingham. 2006. ‘Gravitaxis in Drosophila melanogaster: a forward genetic screen’, *Genes*, Brain and Behavior, 5: 222–39.

Aso, Yoshinori, Divya Sitaraman, Toshiharu Ichinose, Karla R. Kaun, Katrin Vogt, Ghislain Belliart-Guérin, Pierre-Yves Plaçais, Alice A. Robie, Nobuhiro Yamagata, Christopher Schnaitmann, William J. Rowell, Rebecca M. Johnston, Teri-T. B. Ngo, Nan Chen, Wyatt Korff, Michael N. Nitabach, Ulrike Heberlein, Thomas Preat, Kristin M. Branson, Hiromu Tanimoto, and Gerald M. Rubin. 2014. ‘Mushroom body output neurons encode valence and guide memory-based action selection in Drosophila’, eLife, 3: e04580.

Baillargeon, Renée. 1987. ‘Object permanence in 3½- and 4½-month-old infants’, Developmental Psychology, 23: 655–64.

Behbahani, Amir H., Emily H. Palmer, Román A. Corfas, and Michael H. Dickinson. 2021. ‘Drosophila re-zero their path integrator at the center of a fictive food patch’, Current Biology, 31: 4534–46.e5.

Behmer, Spencer T. 2005. ‘Learning in Insects.’ in, Encyclopedia of Entomology (Springer Netherlands: Dordrecht).

Berlyne, D. E. 1950. ‘Novelty and curiosity as determinants of exploratory behaviour’, British Journal of Psychology. General Section, 41: 68–80.

Billeter, Jean-Christophe, Jade Atallah, Joshua J. Krupp, Jocelyn G. Millar, and Joel D. Levine. 2009. ‘Specialized cells tag sexual and species identity in Drosophila melanogaster’, Nature, 461: 987–91.

Bird, Christopher D., and Nathan J. Emery. 2009. ‘Insightful problem solving and creative tool modification by captive nontool-using rooks’, Proceedings of the National Academy of Sciences, 106: 10370–75.

Brodin, Anders. 2005. ‘Mechanisms of cache retrieval in long-term hoarding birds’, Journal of Ethology, 23: 77–83.

Bülthoff, Heinrich, Karl G Götz, and Manfred Herre. 1982. ‘Recurrent inversion of visual orientation in the walking fly, Drosophila melanogaster’, Journal of comparative physiology, 148: 471–81.

Burghardt, Gordon M. 2005. *The genesis of animal play: Testing the limits* (Boston Review: Cambridge, MA, US).

Cheng, K. Y., R. A. Colbath, and M. A. Frye. 2019. ‘Olfactory and Neuromodulatory Signals Reverse Visual Object Avoidance to Approach in Drosophila’, Curr Biol, 29: 2058–65.e2.

Christensen, Janne Winther, Line Peerstrup Ahrendt, Jens Malmkvist, and Christine Nicol. 2021. ‘Exploratory behaviour towards novel objects is associated with enhanced learning in young horses’, Scientific Reports, 11: 1428.

Coen, Philip, Jan Clemens, Andrew J. Weinstein, Diego A. Pacheco, Yi Deng, and Mala Murthy. 2014. ‘Dynamic sensory cues shape song structure in Drosophila’, Nature, 507: 233–37.

Corfas, Román A., Tarun Sharma, and Michael H. Dickinson. 2019. ‘Diverse Food-Sensing Neurons Trigger Idiothetic Local Search in Drosophila’, Current Biology, 29: 1660–68.e4.

Craske, Michelle G., Michael Treanor, Christopher C. Conway, Tomislav Zbozinek, and Bram Vervliet. 2014. ‘Maximizing exposure therapy: An inhibitory learning approach’, Behaviour Research and Therapy, 58: 10–23.

Deutsch, David, Jan Clemens, Stephan Y. Thiberge, Georgia Guan, and Mala Murthy. 2019. ‘Shared Song Detector Neurons in Drosophila Male and Female Brains Drive Sex-Specific Behaviors’, Current Biology, 29: 3200–15.e5.

Diamond, Judy, and Alan Bond. 2003. ‘A comparative analysis of social play in birds’, Behaviour, 140: 1091–115.

Diamond, Mathew E., Moritz von Heimendahl, Per Magne Knutsen, David Kleinfeld, and Ehud Ahissar. 2008. ‘‘Where’ and ‘what’ in the whisker sensorimotor system’, Nature Reviews Neuroscience, 9: 601–12.

Dombrovski, Mark, Martin Y. Peek, Jin-Yong Park, Andrea Vaccari, Marissa Sumathipala, Carmen Morrow, Patrick Breads, Arthur Zhao, Yerbol Z. Kurmangaliyev, Piero Sanfilippo, Aadil Rehan, Jason Polsky, Shada Alghailani, Emily Tenshaw, Shigehiro Namiki, S. Lawrence Zipursky, and Gwyneth M. Card. 2023. ‘Synaptic gradients transform object location to action’, Nature, 613: 534–42.

Dorkenwald, Sven, Arie Matsliah, Amy R. Sterling, Philipp Schlegel, Szi-chieh Yu, Claire E. McKellar, Albert Lin, Marta Costa, Katharina Eichler, Yijie Yin, Will Silversmith, Casey Schneider-Mizell, Chris S. Jordan, Derrick Brittain, Akhilesh Halageri, Kai Kuehner, Oluwaseun Ogedengbe, Ryan Morey, Jay Gager, Krzysztof Kruk, Eric Perlman, Runzhe Yang, David Deutsch, Doug Bland, Marissa Sorek, Ran Lu, Thomas Macrina, Kisuk Lee, J. Alexander Bae, Shang Mu, Barak Nehoran, Eric Mitchell, Sergiy Popovych, Jingpeng Wu, Zhen Jia, Manuel A. Castro, Nico Kemnitz, Dodam Ih, Alexander Shakeel Bates, Nils Eckstein, Jan Funke, Forrest Collman, Davi D. Bock, Gregory S. X. E. Jefferis, H. Sebastian Seung, Mala Murthy, Zairene Lenizo, Austin T. Burke, Kyle Patrick Willie, Nikitas Serafetinidis, Nashra Hadjerol, Ryan Willie, Ben Silverman, John Anthony Ocho, Joshua Bañez, Rey Adrian Candilada, Anne Kristiansen, Nelsie Panes, Arti Yadav, Remer Tancontian, Shirleyjoy Serona, Jet Ivan Dolorosa, Kendrick Joules Vinson, Dustin Garner, Regine Salem, Ariel Dagohoy, Jaime Skelton, Mendell Lopez, Laia Serratosa Capdevila, Griffin Badalamente, Thomas Stocks, Anjali Pandey, Darrel Jay Akiatan, James Hebditch, Celia David, Dharini Sapkal, Shaina Mae Monungolh, Varun Sane, Mark Lloyd Pielago, Miguel Albero, Jacquilyn Laude, Márcia dos Santos, Zeba Vohra, Kaiyu Wang, Allien Mae Gogo, Emil Kind, Alvin Josh Mandahay, Chereb Martinez, John David Asis, Chitra Nair, Dhwani Patel, Marchan Manaytay, Imaan F. M. Tamimi, Clyde Angelo Lim, Philip Lenard Ampo, Michelle Darapan Pantujan, Alexandre Javier, Daril Bautista, Rashmita Rana, Jansen Seguido, Bhargavi Parmar, John Clyde Saguimpa, Merlin Moore, Markus William Pleijzier, Mark Larson, Joseph Hsu, Itisha Joshi, Dhara Kakadiya, Amalia Braun, Cathy Pilapil, Marina Gkantia, Kaushik Parmar, Quinn Vanderbeck, Irene Salgarella, Christopher Dunne, Eva Munnelly, Chan Hyuk Kang, Lena Lörsch, Jinmook Lee, Lucia Kmecova, Gizem Sancer, Christa Baker, Jenna Joroff, Steven Calle, Yashvi Patel, Olivia Sato, Siqi Fang, Janice Salocot, Farzaan Salman, Sebastian Molina-Obando, Paul Brooks, Mai Bui, Matthew Lichtenberger, Edward Tamboboy, Katie Molloy, Alexis E. Santana-Cruz, Anthony Hernandez, Seongbong Yu, Arzoo Diwan, Monika Patel, Travis R. Aiken, Sarah Morejohn, Sanna Koskela, Tansy Yang, Daniel Lehmann, Jonas Chojetzki, Sangeeta Sisodiya, Selden Koolman, Philip K. Shiu, Sky Cho, Annika Bast, Brian Reicher, Marlon Blanquart, Lucy Houghton, Hyungjun Choi, Maria Ioannidou, Matt Collie, Joanna Eckhardt, Benjamin Gorko, Li Guo, Zhihao Zheng, Alisa Poh, Marina Lin, István Taisz, Wes Murfin, Álvaro Sanz Díez, Nils Reinhard, Peter Gibb, Nidhi Patel, Sandeep Kumar, Minsik Yun, Megan Wang, Devon Jones, Lucas Encarnacion-Rivera, Annalena Oswald, Akanksha Jadia, Mert Erginkaya, Nik Drummond, Leonie Walter, Ibrahim Tastekin, Xin Zhong, Yuta Mabuchi, Fernando J. Figueroa Santiago, Urja Verma, Nick Byrne, Edda Kunze, Thomas Crahan, Ryan Margossian, Haein Kim, Iliyan Georgiev, Fabianna Szorenyi, Atsuko Adachi, Benjamin Bargeron, Tomke Stürner, Damian Demarest, Burak Gür, Andrea N. Becker, Robert Turnbull, Ashley Morren, Andrea Sandoval, Anthony Moreno-Sanchez, Diego A. Pacheco, Eleni Samara, Haley Croke, Alexander Thomson, Connor Laughland, Suchetana B. Dutta, Paula Guiomar Alarcón de Antón, Binglin Huang, Patricia Pujols, Isabel Haber, Amanda González-Segarra, Daniel T. Choe, Veronika Lukyanova, Nino Mancini, Zequan Liu, Tatsuo Okubo, Miriam A. Flynn, Gianna Vitelli, Meghan Laturney, Feng Li, Shuo Cao, Carolina Manyari-Diaz, Hyunsoo Yim, Anh Duc Le, Kate Maier, Seungyun Yu, Yeonju Nam, Daniel Bąba, Amanda Abusaif, Audrey Francis, Jesse Gayk, Sommer S. Huntress, Raquel Barajas, Mindy Kim, Xinyue Cui, Gabriella R. Sterne, Anna Li, Keehyun Park, Georgia Dempsey, Alan Mathew, Jinseong Kim, Taewan Kim, Guan-ting Wu, Serene Dhawan, Margarida Brotas, Cheng-hao Zhang, Shanice Bailey, Alexander Del Toro, Runzhe Yang, Stephan Gerhard, Andrew Champion, David J. Anderson, Rudy Behnia, Salil S. Bidaye, Alexander Borst, Eugenia Chiappe, Kenneth J. Colodner, Andrew Dacks, Barry Dickson, Denise Garcia, Stefanie Hampel, Volker Hartenstein, Bassem Hassan, Charlotte Helfrich-Forster, Wolf Huetteroth, Jinseop Kim, Sung Soo Kim, Young-Joon Kim, Jae Young Kwon, Wei-Chung Lee, Gerit A. Linneweber, Gaby Maimon, Richard Mann, Stéphane Noselli, Michael Pankratz, Lucia Prieto-Godino, Jenny Read, Michael Reiser, Katie von Reyn, Carlos Ribeiro, Kristin Scott, Andrew M. Seeds, Mareike Selcho, Marion Silies, Julie Simpson, Scott Waddell, Mathias F. Wernet, Rachel I. Wilson, Fred W. Wolf, Zepeng Yao, Nilay Yapici, Meet Zandawala, and Consortium The FlyWire. 2024. ‘Neuronal wiring diagram of an adult brain’, Nature, 634: 124–38.

Dukas, Reuven. 2008. ‘Evolutionary Biology of Insect Learning’, Annual Review of Entomology, 53: 145–60.

Dukas, Reuven. 2020. ‘Natural history of social and sexual behavior in fruit flies’, Scientific Reports, 10: 21932.

Fayet, Annette L., Erpur Snær Hansen, and Dora Biro. 2020. ‘Evidence of tool use in a seabird’, Proceedings of the National Academy of Sciences, 117: 1277–79.

Galpayage Dona, Hiruni Samadi, Cwyn Solvi, Amelia Kowalewska, Kaarle Mäkelä, HaDi MaBouDi, and Lars Chittka. 2022. ‘Do bumble bees play?’, Animal Behaviour, 194: 239–51.

Glickman, Stephen E., and Richard W. Sroges. 1966. ‘Curiosity in Zoo Animals’, Behaviour, 26: 151–87.

Hindmarsh Sten, Tom, Rufei Li, Florian Hollunder, Shade Eleazer, and Vanessa Ruta. 2025. ‘Male-male interactions shape mate selection in Drosophila’, Cell, 188: 1486–503.e25.

Hoopfer, Eric D., Yonil Jung, Hidehiko K. Inagaki, Gerald M. Rubin, and David J. Anderson. 2015. ‘P1 interneurons promote a persistent internal state that enhances inter-male aggression in Drosophila’, eLife, 4: e11346.

Hulse, Brad K., Hannah Haberkern, Romain Franconville, Daniel Turner-Evans, Shin-ya Takemura, Tanya Wolff, Marcella Noorman, Marisa Dreher, Chuntao Dan, Ruchi Parekh, Ann M. Hermundstad, Gerald M. Rubin, and Vivek Jayaraman. 2021. ‘A connectome of the Drosophila central complex reveals network motifs suitable for flexible navigation and context-dependent action selection’, eLife, 10: e66039.

Ibañez, Victor, Laurens Bohlen, Francesca Manuella, Isabelle Mansuy, Fritjof Helmchen, and Anna-Sophia Wahl. 2023. ‘EXPLORE: a novel deep learning-based analysis method for exploration behaviour in object recognition tests’, Scientific Reports, 13: 4249.

Iwasaki, Kenichi, Charles Neuhauser, Chris Stokes, and Aleksandr Rayshubskiy. 2025. ‘The fruit fly, Drosophila melanogaster, as a microrobotics platform’, Proceedings of the National Academy of Sciences, 122: e2426180122.

Jiang, Lifen, Yaxin Cheng, Shan Gao, Yincheng Zhong, Chengrui Ma, Tianyu Wang, and Yan Zhu. 2020. ‘Emergence of social cluster by collective pairwise encounters in Drosophila’, eLife, 9: e51921.

Kamikouchi, Azusa, Hidehiko K. Inagaki, Thomas Effertz, Oliver Hendrich, André Fiala, Martin C. Göpfert, and Kei Ito. 2009. ‘The neural basis of Drosophila gravity-sensing and hearing’, Nature, 458: 165–71.

Kaplan, Gisela. 2024. ‘The evolution of social play in songbirds, parrots and cockatoos - emotional or highly complex cognitive behaviour or both?’, Neuroscience & Biobehavioral Reviews, 161: 105621.

Kelley, Laura A., and John A. Endler. 2012. ‘Male great bowerbirds create forced perspective illusions with consistently different individual quality’, Proceedings of the National Academy of Sciences, 109: 20980–85.

Kidd, Celeste, and Benjamin Y Hayden. 2015. ‘The Psychology and Neuroscience of Curiosity’, Neuron, 88: 449–60.

Kim, Irene S., and Michael H. Dickinson. 2017. ‘Idiothetic Path Integration in the Fruit Fly Drosophila melanogaster’, Current Biology, 27: 2227–38.e3.

Kladt, Nikolay, and Michael B. Reiser. 2023. ‘Drosophila antennae are dispensable for gravity orientation’, bioRxiv: 2023.03.08.531317.

Kriete, Alexis Lillian, and Karen L. Hollis. 2022. ’Learning in Insects: Perspectives and Possibilities.’ in Mark A. Krause, Karen L. Hollis and Mauricio R. Papini (eds.), Evolution of Learning and Memory Mechanisms (Cambridge University Press: Cambridge).

Kutschireiter, Anna, Melanie A. Basnak, Rachel I. Wilson, and Jan Drugowitsch. 2023. ‘Bayesian inference in ring attractor networks’, Proceedings of the National Academy of Sciences, 120: e2210622120.

Lambert, Megan L., Martina Schiestl, Raoul Schwing, Alex H. Taylor, Gyula K. Gajdon, Katie E. Slocombe, and Amanda M. Seed. 2017. ‘Function and flexibility of object exploration in kea and New Caledonian crows’, Royal Society Open Science, 4: 170652.

Lambert, Poppy J., Alexandra Stiegler, Theresa Rössler, Megan L. Lambert, and Alice M. I. Auersperg. 2021. ’Goffin’s cockatoos discriminate objects based on weight alone’, Biology Letters, 17: 20210250.

Laumer, I. B., T. Bugnyar, S. A. Reber, and A. M. I. Auersperg. 2017. ‘Can hook-bending be let off the hook? Bending/unbending of pliant tools by cockatoos’, Proceedings of the Royal Society B: Biological Sciences, 284: 20171026.

Lee, Su-Yee J., Chris J. Dallmann, Andrew Cook, John C. Tuthill, and Sweta Agrawal. 2025. ‘Divergent neural circuits for proprioceptive and exteroceptive sensing of the Drosophila leg’, Nature Communications, 16: 4105.

Li, Feng, Jack W. Lindsey, Elizabeth C. Marin, Nils Otto, Marisa Dreher, Georgia Dempsey, Ildiko Stark, Alexander S. Bates, Markus William Pleijzier, Philipp Schlegel, Aljoscha Nern, Shin-ya Takemura, Nils Eckstein, Tansy Yang, Audrey Francis, Amalia Braun, Ruchi Parekh, Marta Costa, Louis K. Scheffer, Yoshinori Aso, Gregory S. X. E. Jefferis, Larry F. Abbott, Ashok Litwin-Kumar, Scott Waddell, and Gerald M. Rubin. 2020. ‘The connectome of the adult Drosophila mushroom body provides insights into function’, eLife, 9: e62576.

Liu, Q., K. Simpson, P. Izar, E. Ottoni, E. Visalberghi, and D. Fragaszy. 2009. ‘Kinematics and energetics of nut-cracking in wild capuchin monkeys (Cebus libidinosus) in Piauí, Brazil’, American journal of physical anthropology, 138: 210–20.

Lobo, Michele A., Elena Kokkoni, Ana Carolina de Campos, and James C. Galloway. 2014. ‘Not just playing around: Infants’ behaviors with objects reflect ability, constraints, and object properties’, Infant Behavior and Development, 37: 334–51.

Loukola, Olli J., Cwyn Solvi, Louie Coscos, and Lars Chittka. 2017. ‘Bumblebees show cognitive flexibility by improving on an observed complex behavior’, Science, 355: 833–36.

Lyu, Cheng, L. F. Abbott, and Gaby Maimon. 2022. ‘Building an allocentric travelling direction signal via vector computation’, Nature, 601: 92–97.

Maimon, Gaby, Andrew D. Straw, and Michael H. Dickinson. 2008. ‘A Simple Vision-Based Algorithm for Decision Making in Flying Drosophila’, Current Biology, 18: 464–70.

Mather, Jennifer A., and Roland C. Anderson. 1999. ‘Exploration, play and habituation in octopuses (Octopus dofleini)’, Journal of Comparative Psychology, 113: 333–38.

Matsliah, Arie, Amy Sterling, Sven Dorkenwald, Kai Kuehner, Ryan Morey, Hyunjune Seung, and Mala Murthy. 2023. Codex: Connectome Data Explorer.

McCoy, Dakota E., Martina Schiestl, Patrick Neilands, Rebecca Hassall, Russell D. Gray, and Alex H. Taylor. 2019. ‘New Caledonian Crows Behave Optimistically after Using Tools’, Current Biology, 29: 2737–42.e3.

Mizuuchi, Ryo, Hiroshi Kawase, Hirofumi Shin, Daisuke Iwai, and Shigeru Kondo. 2018. ‘Simple rules for construction of a geometric nest structure by pufferfish’, Scientific Reports, 8: 12366.

Mongeau, Jean-Michel, Karen Y. Cheng, Jacob Aptekar, and Mark A. Frye. 2019. ‘Visuomotor strategies for object approach and aversion in Drosophila melanogaster’, Journal of Experimental Biology, 222.

Mussells Pires, Peter, Lingwei Zhang, Victoria Parache, L. F. Abbott, and Gaby Maimon. 2024. ‘Converting an allocentric goal into an egocentric steering signal’, Nature, 626: 808–18.

Ofstad, Tyler A., Charles S. Zuker, and Michael B. Reiser. 2011. ‘Visual place learning in Drosophila melanogaster’, Nature, 474: 204–07.

Ogasawara, Takaya, Fatih Sogukpinar, Kaining Zhang, Yang-Yang Feng, Julia Pai, Ahmad Jezzini, and Ilya E. Monosov. 2022. ‘A primate temporal cortex–zona incerta pathway for novelty seeking’, Nature Neuroscience, 25: 50–60.

Pepperberg, I. M., M. R. Willner, and L. B. Gravitz. 1997. ‘Development of Piagetian object permanence in a grey parrot (Psittacus erithacus)’, J Comp Psychol, 111: 63–75.

Robie, Alice A., Andrew D. Straw, and Michael H. Dickinson. 2010. ‘Object preference by walking fruit flies, Drosophila melanogaster, is mediated by vision and graviperception’, Journal of Experimental Biology, 213: 2494–506.

Schlegel, Philipp, Yijie Yin, Alexander S. Bates, Sven Dorkenwald, Katharina Eichler, Paul Brooks, Daniel S. Han, Marina Gkantia, Marcia dos Santos, Eva J. Munnelly, Griffin Badalamente, Laia Serratosa Capdevila, Varun A. Sane, Alexandra M. C. Fragniere, Ladann Kiassat, Markus W. Pleijzier, Tomke Stürner, Imaan F. M. Tamimi, Christopher R. Dunne, Irene Salgarella, Alexandre Javier, Siqi Fang, Eric Perlman, Tom Kazimiers, Sridhar R. Jagannathan, Arie Matsliah, Amy R. Sterling, Szi-chieh Yu, Claire E. McKellar, Krzysztof Kruk, Doug Bland, Zairene Lenizo, Austin T. Burke, Kyle Patrick Willie, Alexander S. Bates, Nikitas Serafetinidis, Nashra Hadjerol, Ryan Willie, Ben Silverman, John Anthony Ocho, Joshua Bañez, Rey Adrian Candilada, Jay Gager, Anne Kristiansen, Nelsie Panes, Arti Yadav, Remer Tancontian, Shirleyjoy Serona, Jet Ivan Dolorosa, Kendrick Joules Vinson, Dustin Garner, Regine Salem, Ariel Dagohoy, Jaime Skelton, Mendell Lopez, Thomas Stocks, Anjali Pandey, Darrel Jay Akiatan, James Hebditch, Celia David, Dharini Sapkal, Shaina Mae Monungolh, Varun Sane, Mark Lloyd Pielago, Miguel Albero, Jacquilyn Laude, Márcia dos Santos, David Deutsch, Zeba Vohra, Kaiyu Wang, Allien Mae Gogo, Emil Kind, Alvin Josh Mandahay, Chereb Martinez, John David Asis, Chitra Nair, Dhwani Patel, Marchan Manaytay, Clyde Angelo Lim, Philip Lenard Ampo, Michelle Darapan Pantujan, Daril Bautista, Rashmita Rana, Jansen Seguido, Bhargavi Parmar, John Clyde Saguimpa, Merlin Moore, Markus W. Pleijzier, Mark Larson, Joseph Hsu, Itisha Joshi, Dhara Kakadiya, Amalia Braun, Cathy Pilapil, Kaushik Parmar, Quinn Vanderbeck, Christopher Dunne, Eva Munnelly, Chan Hyuk Kang, Lena Lörsch, Jinmook Lee, Lucia Kmecova, Gizem Sancer, Christa Baker, Jenna Joroff, Steven Calle, Yashvi Patel, Olivia Sato, Janice Salocot, Farzaan Salman, Sebastian Molina-Obando, Mai Bui, Matthew Lichtenberger, Edmark Tamboboy, Katie Molloy, Alexis E. Santana-Cruz, Anthony Hernandez, Seongbong Yu, Marissa Sorek, Arzoo Diwan, Monika Patel, Travis R. Aiken, Sarah Morejohn, Sanna Koskela, Tansy Yang, Daniel Lehmann, Jonas Chojetzki, Sangeeta Sisodiya, Selden Koolman, Philip K. Shiu, Sky Cho, Annika Bast, Brian Reicher, Marlon Blanquart, Lucy Houghton, Hyungjun Choi, Maria Ioannidou, Matt Collie, Joanna Eckhardt, Benjamin Gorko, Li Guo, Zhihao Zheng, Alisa Poh, Marina Lin, István Taisz, Wes Murfin, Álvaro Sanz Díez, Nils Reinhard, Peter Gibb, Nidhi Patel, Sandeep Kumar, Minsik Yun, Megan Wang, Devon Jones, Lucas Encarnacion-Rivera, Annalena Oswald, Akanksha Jadia, Mert Erginkaya, Nik Drummond, Leonie Walter, Ibrahim Tastekin, Xin Zhong, Yuta Mabuchi, Fernando J. Figueroa Santiago, Urja Verma, Nick Byrne, Edda Kunze, Thomas Crahan, Ryan Margossian, Haein Kim, Iliyan Georgiev, Fabianna Szorenyi, Atsuko Adachi, Benjamin Bargeron, Tomke Stürner, Damian Demarest, Burak Gür, Andrea N. Becker, Robert Turnbull, Ashley Morren, Andrea Sandoval, Anthony Moreno-Sanchez, Diego A. Pacheco, Eleni Samara, Haley Croke, Alexander Thomson, Connor Laughland, Suchetana B. Dutta, Paula Guiomar Alarcón de Antón, Binglin Huang, Patricia Pujols, Isabel Haber, Amanda González-Segarra, Albert Lin, Daniel T. Choe, Veronika Lukyanova, Nino Mancini, Zequan Liu, Tatsuo Okubo, Miriam A. Flynn, Gianna Vitelli, Meghan Laturney, Feng Li, Shuo Cao, Carolina Manyari-Diaz, Hyunsoo Yim, Anh Duc Le, Kate Maier, Seungyun Yu, Yeonju Nam, Daniel Bąba, Amanda Abusaif, Audrey Francis, Jesse Gayk, Sommer S. Huntress, Raquel Barajas, Mindy Kim, Xinyue Cui, Amy R. Sterling, Gabriella R. Sterne, Anna Li, Keehyun Park, Georgia Dempsey, Alan Mathew, Jinseong Kim, Taewan Kim, Guan-ting Wu, Serene Dhawan, Margarida Brotas, Cheng-hao Zhang, Shanice Bailey, Alexander Del Toro, Kisuk Lee, Thomas Macrina, Casey Schneider-Mizell, Sergiy Popovych, Oluwaseun Ogedengbe, Runzhe Yang, Akhilesh Halageri, Will Silversmith, Stephan Gerhard, Andrew Champion, Nils Eckstein, Dodam Ih, Nico Kemnitz, Manuel Castro, Zhen Jia, Jingpeng Wu, Eric Mitchell, Barak Nehoran, Shang Mu, J. Alexander Bae, Ran Lu, Ryan Morey, Kai Kuehner, Derrick Brittain, Chris S. Jordan, David J. Anderson, Rudy Behnia, Salil S. Bidaye, Alexander Borst, Eugenia Chiappe, Forrest Collman, Kenneth J. Colodner, Andrew Dacks, Barry Dickson, Jan Funke, Denise Garcia, Stefanie Hampel, Volker Hartenstein, Bassem Hassan, Charlotte Helfrich-Forster, Wolf Huetteroth, Jinseop Kim, Sung Soo Kim, Young-Joon Kim, Jae Young Kwon, Wei-Chung Lee, Gerit A. Linneweber, Gaby Maimon, Richard Mann, Stéphane Noselli, Michael Pankratz, Lucia Prieto-Godino, Jenny Read, Michael Reiser, Katie von Reyn, Carlos Ribeiro, Kristin Scott, Andrew M. Seeds, Mareike Selcho, Marion Silies, Julie Simpson, Scott Waddell, Mathias F. Wernet, Rachel I. Wilson, Fred W. Wolf, Zepeng Yao, Nilay Yapici, and Consortium FlyWire. 2024. ‘Whole-brain annotation and multi-connectome cell typing of Drosophila’, Nature, 634: 139–52.

Schrauf, Cornelia, Ludwig Huber, and Elisabetta Visalberghi. 2008. ‘Do capuchin monkeys use weight to select hammer tools?’, Animal Cognition, 11: 413–22.

Schretter, Catherine E., Yoshinori Aso, Alice A. Robie, Marisa Dreher, Michael-John Dolan, Nan Chen, Masayoshi Ito, Tansy Yang, Ruchi Parekh, Kristin M. Branson, and Gerald M. Rubin. 2020. ‘Cell types and neuronal circuitry underlying female aggression in Drosophila’, eLife, 9: e58942.

Seelig, Johannes D., and Vivek Jayaraman. 2015. ‘Neural dynamics for landmark orientation and angular path integration’, Nature, 521: 186–91.

Shaw, Paul J., Chiara Cirelli, Ralph J. Greenspan, and Giulio Tononi. 2000. ‘Correlates of Sleep and Waking in Drosophila melanogaster’, Science, 287: 1834–37.

Sun, Yuanjie, Rong Qiu, Xiaonan Li, Yaxin Cheng, Shan Gao, Fanchen Kong, Li Liu, and Yan Zhu. 2020. ‘Social attraction in Drosophila is regulated by the mushroom body and serotonergic system’, Nature Communications, 11: 5350.

Triphan, Tilman, Clara H. Ferreira, and Wolf Huetteroth. 2025. ‘Play-like behavior exhibited by the vinegar fly Drosophila melanogaster’, Current Biology, 35: 1145–55.e2.

Turner-Evans, Daniel, Stephanie Wegener, Hervé Rouault, Romain Franconville, Tanya Wolff, Johannes D. Seelig, Shaul Druckmann, and Vivek Jayaraman. 2017. ‘Angular velocity integration in a fly heading circuit’, eLife, 6: e23496.

Tuthill, John C, and Rachel I Wilson. 2016. ‘Mechanosensation and Adaptive Motor Control in Insects’, Current Biology, 26: R1022–R38.

von Reyn, Catherine R., Patrick Breads, Martin Y. Peek, Grace Zhiyu Zheng, W. Ryan Williamson, Alyson L. Yee, Anthony Leonardo, and Gwyneth M. Card. 2014. ‘A spike-timing mechanism for action selection’, Nature Neuroscience, 17: 962–70.

Wang, Liming, Xiaoqing Han, Jennifer Mehren, Makoto Hiroi, Jean-Christophe Billeter, Tetsuya Miyamoto, Hubert Amrein, Joel D. Levine, and David J. Anderson. 2011. ‘Hierarchical chemosensory regulation of male-male social interactions in Drosophila’, Nature Neuroscience, 14: 757–62.

Wehner, R. 1972. ‘Spontaneous pattern preferences of Drosophila melanogaster to black areas in various parts of the visual field’, Journal of insect physiology, 18: 1531–43.

Wolff, Tanya, Mark Eddison, Nan Chen, Aljoscha Nern, Preeti Sundaramurthi, Divya Sitaraman, and Gerald M. Rubin. 2025. “Cell type-specific driver lines targeting the Drosophila central complex and their use to investigate neuropeptide expression and sleep regulation.” In.: eLife Sciences Publications, Ltd.

Yamamoto, Daisuke, and Masayuki Koganezawa. 2013. ‘Genes and circuits of courtship behaviour in Drosophila males’, Nature Reviews Neuroscience, 14: 681–92.

